# Structure-based design of a chemical probe set for the 5-HT_5A_ serotonin receptor

**DOI:** 10.1101/2021.12.06.471449

**Authors:** Anat Levit Kaplan, Ryan T. Strachan, Joao M. Braz, Veronica Craik, Samuel Slocum, Thomas Mangano, Vanessa Amabo, Henry O’Donnell, Parnian Lak, Allan I. Basbaum, Bryan L. Roth, Brian Shoichet

## Abstract

The 5-HT_5A_ receptor (5-HT_5A_R), for which no selective agonists and only a few antagonists exist, remains the least understood serotonin (5-HT) receptor. A single commercial antagonist (SB-699551) has been widely used to investigate central nervous system (CNS) 5-HT_5A_R function in neurological disorders, including pain. However, because SB-699551 has affinity for many 5-HTRs, lacks inactive property-matched controls, and has assay interference concerns, it has liabilities as a chemical probe. To better illuminate 5-HT_5A_R function, we developed a probe set through iterative rounds of molecular docking, pharmacological testing, and optimization. Docking over six million lead-like molecules against a 5-HT_5A_R homology model identified five mid-μM ligand starting points with unique scaffolds. Over multiple rounds of structure-based design and testing, a new quinoline scaffold with high affinity and enhanced selectivity for the 5-HT_5A_R was developed, leading to UCSF678, a 42 nM arrestin-biased partial agonist at the 5-HT_5A_R with a much more restricted off-target profile and decreased assay liabilities vs. SB-699551. Site-directed mutagenesis supported the docked pose of UCSF678, which was also consistent with recent published 5-HTR structures. Surprisingly, property-matched analogs of UCSF678 that were either inactive across 5-HTRs or retained affinity for UCSF678’s off-targets revealed that 5-HT_5A_R engagement is nonessential for alleviating pain in a mouse model, contrary to previous studies using less-selective ligands. Relative to SB-699551, these molecules constitute a well-characterized and more selective probe set with which to study the function of the 5-HT_5A_ receptor.

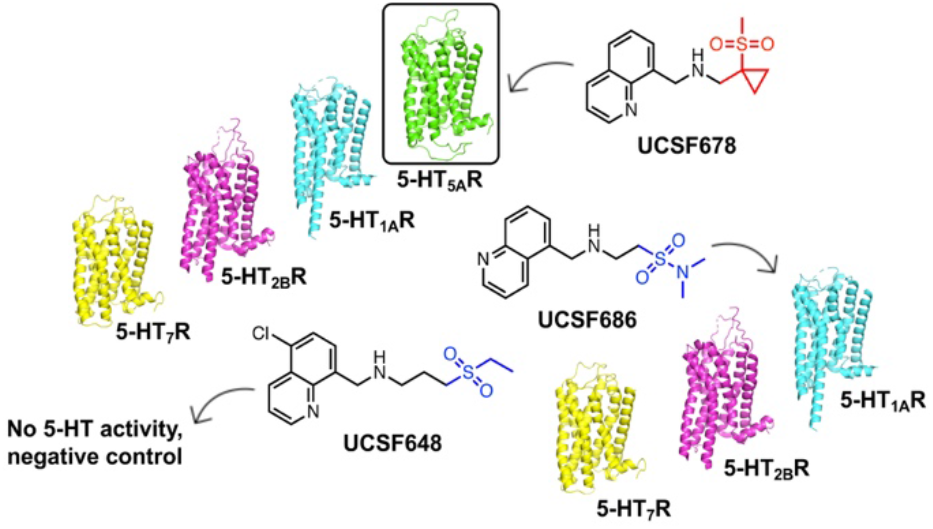
Table of Contents Graphic.

## INTRODUCTION

Though G protein-coupled receptors (GPCRs) constitute a plurality of drug targets, many remain underexploited (*1–3*). Scalable methods to identify small molecule probes to illuminate their (patho)physiological roles include open-source physical assays (*4*), cheminformatic ligand-based virtual screens (*5, 6*), and structure-based virtual screening campaigns (*7*). For these under-studied receptors, homology models have been used to template probe discovery, including recent efforts against the orphan receptors GPR68 and GPR65 (*8*), the primate-exclusive novel opioid receptor MRGPRX2 (*9*), and others (*10, 11*).

The human serotonin receptors (5-HTRs) are prototypic drug targets that modulate key neurological processes, including aggression, anxiety, appetite, cognition, learning, memory, mood, sleep, and thermoregulation (*12*). Of the 14 serotonin receptor subtypes, the 5-HT_5A_R is perhaps the least understood, largely due to the lack of readily-available, selective chemical probes (*13*). First cloned in 1994 (*14*), the human 5-HT_5A_R most closely resembles the 5-HT_1_R in terms of ligand recognition (e.g., preference for binding methiothepin and ergotamine), Gi/o coupling, and presumed inhibitory autoreceptor function in human cortical and hippocampal pyramidal neurons (*15*). 5-HT_5A_Rs are confined primarily to neuronal components of the nervous system and genetic and pharmacological studies have associated them with several neurological functions (reviewed by (*13*), (*15*)). More recent studies also implicate the 5-HT_5A_R in memory stabilization (*16*) and memory deficits associated with forgetting and amnesia (*17*). Consistent with a 5-HT_5A_R contribution to psychiatric disorders, the receptor’s inactivity is required for the efficacy of chronic selective serotonin reuptake inhibitor (SSRI) antidepressant treatment (*18*). Furthermore, dense immunolabelling in the dorsal horn and Onuff’s nucleus of the spinal cord suggests that 5-HT_5A_ receptors modulate central motor control, control of pelvic floor musculature and nociception (*19*). Of note, recent *in vivo* studies demonstrated a 5-HT_5A_R contribution to nociceptive processing in both naïve and injured mice. (*20, 21*).

Notwithstanding its association with these multiple physiological processes, selective probe molecules for the 5-HT_5A_R are unavailable to the community, limiting our ability to test the relevance of these largely genetic associations pharmacologically, or to leverage them for therapeutic development. Currently, only a few molecules have been mooted as even modestly selective antagonists for the receptor (*15, 22*) and selective agonists have yet to be described. Accordingly, much of what we know about 5-HT_5A_R pharmacology and its (dys)function stems from blockade with the commercially available 5-HT_5A_R antagonist SB-699551 (*23*). However, SB-699551 possesses considerable off-target activity (≤1μM) for many 5-HTR family members and lacks a chemically matched negative control probe with which to control for them. Indeed, the off-target activities of SB-699551 and of another 5-HT_5A_R antagonist A-843277, the latter of which is unavailable commercially, were substantial enough to confound interpretation of *in vivo* studies (*24*).

To better illuminate 5-HT_5A_R function, here we deployed an iterative molecular docking and empirical testing strategy, adapted from one successfully used with the understudied receptors GPR65 and GPR68 (*8*), and with MRGPRX2 (*9*). Docking of >6 million lead-like molecules against a homology model of the receptor initiated an iterative cycle of analoging, docking, and pharmacological testing that ultimately identified a novel chemical scaffold with mid-nanomolar affinity for 5-HT_5A_R and a far more restricted off-target profile versus SB-699551. Property-matched probe-pairs that controlled for off-target activities were also developed, and the set of probe molecules was used to investigate a previously hypothesized role for the 5-HT_5A_R in neuropathic pain, here in a mouse model.

## MATERIALS AND METHODS

### Homology modeling

A homology model of the 5-HT_5A_R was calculated using the crystal structure of 5-HT_1B_R in complex with Ergotamine as the template (PDB: 4IAR (chain A), 4IAQ (chain A)(*25*). The sequence of the target, template, and several members of the 5-HTR family were aligned using PROMALS3D (*26*), using sequences of human 5-HT_1B_R (Uniprot accession number: P28222), 5-HT_2A_R (P28223), and 5-HT_2B_R (P41595). The alignment was manually edited to (1) remove 31 residues from the amino terminus of 5-HT_5A_R and one residue from the carboxy terminus that extended past the resolved template structure; (2) remove the engineered apocytochrome b562 RIL (BRIL) from the template and the corresponding residues in ICL3 of 5-HT_5A_R. The final sequence alignment is shown in **Figure 1A**. Based on this alignment, 1000 homology models were built using MODELER-9v15 (*27*). Ergotamine was retained in the modeling to ensure a ligand-competent orthosteric site. The resulting models were evaluated for their ability to enrich known 5-HT_5A_R ligands over property-matched decoys through docking to the orthosteric site, using DOCK 3.7 (*28*) (below). Decoy molecules share the physical properties of known ligands but are topologically distinct from them and so are unlikely to bind, thus controlling for the enrichment of molecules by physical properties alone. For this aim, 17 known 5-HT_5A_R ligands with MW < 450 were extracted from the IUPHAR database (*29*), and 1133 property-matched decoys were generated using the DUD-E server (*30*). The 1000 homology models were ranked by their ability to highly enrich the known ligands over the decoy molecules, using adjusted logAUC (*30*) and the enrichment factor at 1% of the database (EF_1%_), both of which bias for early enrichment, and by the fidelity of the docked pose of ergotamine to the crystallographic structure in the template structure. The best scoring model was further optimized through minimization with the AMBER protein force field and the GAFF ligand force field supplemented with AM1BCC charges (*31*). The integrity of the minimized model was assessed by redocking the known ligands and decoy molecules and re-calculating enrichment factors.

**Figure 1.**
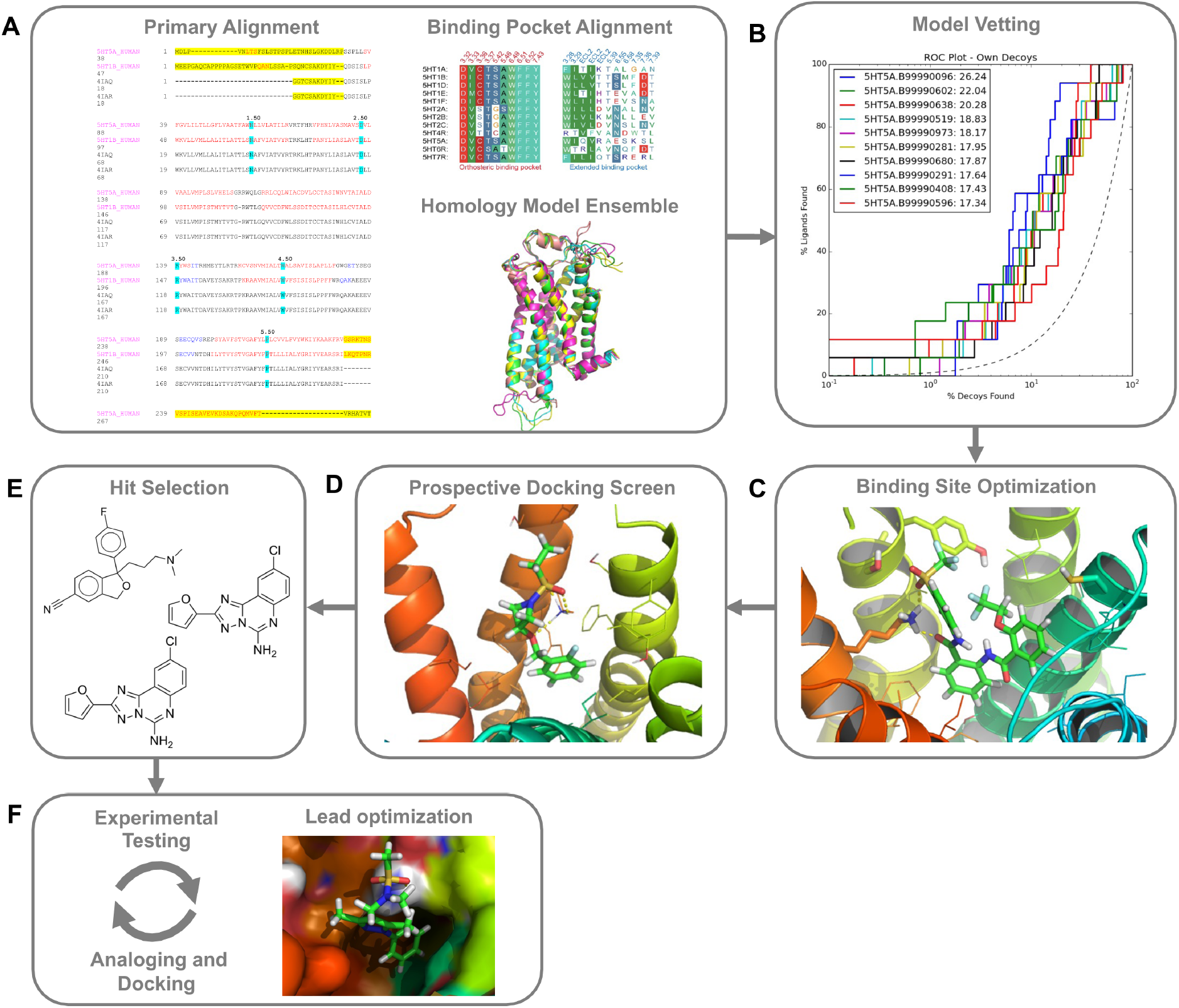
Structure-based strategy for discovery of novel 5-HT5AR chemotypes. **(A)** Iterative docking and empirical testing begins with generation of an ensemble of homology models based on the alignment between 5-HT5AR and 5-HT1BR and relevant crystal structures. Residues highlighted in cyan are the x.50 positions, which are most conserved in each helix. At top right is the alignment of binding site residues for all 5-HT receptors, and their corresponding Ballesteros-Weinstein numbers (*46*). At the panel bottom, an ensemble of 1000 homology models were built using MODELER-9v15, with ergotamine retained in the modeling for a ligand-competent orthosteric site. **(B)** The homology models were then evaluated for their ability to enrich known 5-HT5AR ligands over property-matched decoys through docking to the orthosteric site, using DOCK 3.7 (*28*). Shown is the enrichment curves for the top 10 performing 5-HT5AR models. **(C)** The binding site of the best performing model was further optimized through energy minimization. The performance of the minimized model was assessed by redocking the known ligands and decoy molecules and re-calculating enrichment factors. **(D)** Over 6 million “lead-like” molecules from ZINC15 were prospectively screened against the 5-HT5AR model by molecular docking. **(E)** The top 3,000 scoring molecules were filtered for chemical novelty against ∼28,000 annotated aminergic ligands (Tc < 0.40) and resulting molecules ranked according to favorable geometries and interactions with key binding site residues (e.g., D121^3.32^). (F) Representative compounds were tested in binding assays, and active compounds were optimized for potency and affinity through an cycle of analoging, docking, and testing.

### Virtual ligand screening and selection of potential ligands for experimental testing

The orthosteric site of the 5-HT_5A_R model was prospectively screened against >6 million “lead-like” molecules from the ZINC15 database (http://zinc15.docking.org/) using DOCK3.7 (*28*). DOCK3.7 fits pre-generated flexible ligands into a binding site by superimposing atoms of each molecule on local hot-spots in the site (“matching spheres”), representing favorable positions for individual ligand atoms. Here, 45 matching spheres were used, drawn from the docked pose of lysergic acid diethylamide (LSD). The resulting docked ligand poses were scored by summing the receptor-ligand electrostatics and van der Waals interaction energies and corrected for context-dependent ligand desolvation (*32, 33*). Receptor structures were protonated using Reduce (*34*). Partial charges from the united-atom AMBER (*31*) force field were used for all receptor atoms. Potential energy grids for the different energy terms of the scoring function were pre-calculated based on the AMBER potential (*31*) for the van der Waals term and the Poisson-Boltzmann method QNIFFT (*35, 36*) for electrostatics. Context-dependent ligand desolvation was calculated using an adaptation of the Generalized-Born method (*32*). Ligands were protonated with Marvin (version 15.11.23.0, ChemAxon, 2015; https://www.chemaxon.com), at pH 7.4. Each protomer was rendered into 3D using Corina (v.3.6.0026, Molecular Networks GmbH; https://www.mn-am.com/products/corina), and conformationally sampled using Omega (v.2.5.1.4, OpenEye Scientific Software; https://www.eyesopen.com/omega). Ligand atomic charges and initial desolvation energies were calculated as described (*37*). In the docking screen, each library molecule was sampled in about 16,000 orientations and, on average, 350 conformations. The best scoring configuration for each docked molecule was relaxed by rigid-body minimization. Overall, about 6.11 × 10^12^ complexes were sampled and scored; this took 3008 core hours— spread over 100 cores, about 30 hours of wall-clock time.

By calculating ECFP4-based Tanimoto coefficients (Tc) against ∼28,000 annotated aminergic ligands (acting at dopamine, serotonin and adrenergic receptors), extracted from the ChEMBL20 database (*38*), we filtered the top 3,000 ranked molecules emerging from the screen for topological dissimilarity to known ligands. Molecules with Tc < 0.40 to these aminergic ligands were considered as dissimilar and passed this filter. The remaining molecules were visually inspected in their docked poses. Topologically diverse molecules that adopted favorable geometries and formed specific interactions with binding site residues, such as an ion-pair with D121^3.32^ and hydrogen bonds with residues on extracellular loop 2 (ECL2), were prioritized from among the top 2,000 docking-ranked molecules that remained. Twenty-five compounds were selected for initial experimental testing.

### Hit optimization

Potential analogs of the hit compound 5A-6 (ZINC000089807724) were identified in five iterative rounds through a combination of similarity and sub-structure searches of the ZINC database (*37*). In each iteration, analogs were docked to the 5-HT_5A_R orthosteric site using DOCK3.7. As was true in the primary screen, the resulting docked poses were manually evaluated for specific interactions and compatibility with the site, and prioritized analogs were acquired and tested experimentally.

### Compound handling

Manually curated hits were purchased from Enamine, Chembridge, and Molport. Portions of each compound (0.5 to 1 mg) were dissolved in DMSO at 10mM and maintained as −20°C stock solutions. Freeze-thaw cycles were minimized, and new stocks of compounds were made from the original dry stocks for additional rounds of binding and activity profiling.

### Molecular biology and site-directed mutagenesis

An in-frame fusion between the human 5-HT_5A_R from the Presto-Tango cDNA library (*4*) and the human G*α*i1 was made via HiFi DNA assembly (New England Biolabs, Ipswich, MA). Mutations of key contact points between docked ligands and the human 5-HT_5A_R binding pocket were made via site-directed mutagenesis as directed (Stratagene, La Jolla, CA). Quickchange II mutagenic primer sets were the following. W117A 5-HT_5A_R: 5’aaagtacgtcacatgcaatcgccaactgacaaagtcttcgtc-3’ and 5’- gacgaagactttgtcagttggcgattgcatgtgacgtacttt-3’. Q193A 5-HT_5A_R: 5’- ggctcccgactgaccgcgcattcctctgatcc-3’, 5’-ggatcagaggaatgcgcggtcagtcgggagcc-3’. Q193L 5-HT5A: 5’-aaggctcccgactgacaaggcattcctctgatcc-3’, 5’-ggatcagaggaatgccttgtcagtcgggagcctt-3’. Q193F 5- HT_5A_R: 5’-gaaggctcccgactgacgaagcattcctctgatccct-3’, 5’- agggatcagaggaatgcttcgtcagtcgggagccttc-3’. Individual clones were selected and sequence-verified (GeneWiz, Morrisville, NC).

### Cell culture

To generate membranes expressing high amounts of receptor, suspension Expi293F cells were cultured and transfected exactly as stated by the manufacturer (ThermoFisher Scientific, Waltham, MA). Briefly, Expi293F cells were maintained in vented 125mL polycarbonate Erlenmeyer flasks (GeneMate, Radnor, PA) in 30mL of Expi293 growth medium at 37°C, 8% CO2, and 115 rpm. For BRET2 functional studies, HEK293T cells obtained from ATCC (Manassas, VA) were maintained in DMEM containing 10% FBS, 100 Units/mL penicillin, and 100 μg/mL streptomycin (Gibco-ThermoFisher, Waltham, MA) in a humidified atmosphere at 37°C and 5% CO2. For transfection and BRET2 assays, HEKT cells were split into DMEM containing 1% dialyzed FBS, 100 Units/mL penicillin, and 100 μg/mL streptomycin (see ‘BRET2 functional assays’ below) to minimize exposure to 5-HT in serum. For Tango assays that also include GPCRome screens, HTLA cells (a HEK293 cell line stably expressing the tTA-dependent luciferase reporter and the β-arrestin2-TEV fusion protein, (a gift from the laboratory of R. Axel) were maintained in DMEM supplemented with 10% FBS, 100 U/ml penicillin, 100 μg/ml streptomycin, 2.0 μg/ml puromycin, and 100 μg/ml hygromycin B in a humidified atmosphere at 37°C and 5% CO2. For transfection and Tango assays, HTLA cells were cultured in DMEM containing 1% dialyzed FBS, 100 Units/mL penicillin, and 100 μg/mL streptomycin.

### BRET2 functional assays

Cells were plated in six-well dishes at a density of 700,000 to 800,000 cells/well or in 10-cm dishes at a density of 7 to 8 million cells/dish. Cells were transfected 2 to 4 hours later using a 1:1:1:1 DNA ratio of receptor:Gα-RLuc8:Gβ:Gγ-GFP2 (Olsen et al. NatChemBiol 2020). DNA amounts were 100ng and 750ng per construct for 6-well and 10cm dishes, respectively. Transit 2020 (Mirus Biosciences, Madison, WI) was used to complex the DNA at a ratio of 3.0 μL Transit/μg DNA in OptiMEM (10 ng DNA/µL OptiMEM, Gibco-ThermoFisher, Waltham, MA). The next day, cells were harvested from the plate using Versene solution (phosphate buffered saline buffer + 0.5 mM EDTA, pH 7.4), and plated in poly-L-lysine-coated white wall, clear bottom 96-well assay plates (Greiner Bio-One, Monroe, NC) at a density of 30,000-50,000 cells/well. One day after plating, white backings (Perkin Elmer, Waltham, MA) were applied to the plate bottoms, and growth medium was carefully aspirated and replaced with 60 μL of assay buffer (1x HBSS + 20 mM HEPES, pH 7.4). Ten microliters of freshly prepared 50 μM Coelenterazine 400a (Nanolight Technologies, Pinetop, AZ) were added to each well; 5 min later cells were treated with 30 μL of drug. Plates were read 5 min later on an LB940 Mithras plate reader (Berthold Technologies, Oak Ridge, TN) with 395 nm (RLuc8-coelenterazine 400a) and 510 nm (GFP2) emission filters, at 1 second/well integration times. Plates were read serially six times and stable measurements from the fourth read were used in all analyses. The BRET2 ratio was computed as the ratio of GFP2 emission to RLuc8 emission.

### Tango β-arrestin2 recruitment assay

For analog screening, HTLA cells were plated on Day 1 at a density of 10 × 10^6^ cells per 150 mm cell-culture dish and transfected with 20μg Tango receptor cDNA (*39*) via the calcium phosphate method the following day (Day 2) (*40*) (*4*). Twenty-four hours post-transfection (Day 3), cells were plated into white-wall, clear bottom 384-well plates (Greiner Bio-One, Monroe, NC) at 15,000 cells per well in 40μL DMEM containing 1% dialyzed FBS, 100 Units/mL penicillin, and 100 μg/mL streptomycin. Twenty-four hours later (Day 4), cells were treated with 20μL of a single maximal concentration (10μM) or a ligand serial dilution in assay buffer (20mM HEPES, 1X HBSS, 0.1% fatty acid free BSA, 0.01% ascorbic acid, pH7.4). Approximately 18-20 hrs later (Day 5), 20μL of a 1/5^th^ diluted solution of Bright-Glo reagent (Promega, Madison, WI) was added directly to the wells, incubated for 10min at RT, light adapted for 30sec, and read for 0.5sec per well in a Spectramax luminescence plate reader (Molecular Devices, San Jose, CA).

GPCRome-wide screening was accomplished using previously described methods with several modifications (*4*). First, HTLA cells were plated in DMEM containing 2% dialyzed FBS, 100 Units/mL penicillin, and 100 μg/mL streptomycin. Next, the cells were transfected using an in-plate polyethylenimine (PEI) method (*41*). Tango receptor DNAs were resuspended in OptiMEM and hybridized with PEI prior to dilution and distribution into 384-well plates and subsequent addition to cells. After overnight incubation, drugs diluted in DMEM with 1% dialyzed FBS were added to cells without replacement of the medium. Approximately 18-20 hrs later, luciferin substrate was added, and luminescence measured as detailed above.

### Membrane preparation

Membranes for radioligand binding and GTP*γ*[^35^S] loading experiments were prepared via differential centrifugation as follows. Expi293F suspension cells were split at a density of 75 x 10^6^ cells in 25.5mL of growth medium and transfected with a DNA complexation mixture containing 30 μg of 5-HT_5A_R (for binding) or 5-HT_5A_R-Gi1 fusion cDNA (for GTP*γ*[^35^S] loading), 80 μL of Expifectamine, and 3mL of OptiMEM. Approximately 18 hr post-transfection, Enhancers 1 and 2 were added. Cells were harvested 48 hr post-transfection via centrifugation at 200 x g for 10min at 4°C. The cell pellet was resuspended in 10mL of homogenization buffer (50 mM TrisHCl, 2mM EDTA and protease inhibitors 500 μM AEBSF, 1.0 μM E-64, 1.0 μM leupeptin, 150 nM aprotinin; pH7.4) and dounce homogenized on ice. Cell debris was removed at 500 x g for 10min at 4°C, and microsomes were recovered from the low speed supernatant at 35,000 x g for 60min at 4°C. The high-speed pellet was resuspended in 0.5-1.0 mL resuspension buffer (50mM TrisHCl, 2mM EDTA, 10mM MgCl_2_, 5% glycerol, and protease inhibitors 500 μM AEBSF, 1.0 μM E-64, 1μM leupeptin, 150nM aprotinin; pH7.4) and aliquoted into 1.5 mL tubes. For GTP*γ*[^35^S] loading assays, the microsome suspension was immediately frozen and stored at −80°C. For binding, microsomes were recovered at 16,000 x g for 15min at 4°C, followed by removal of the supernatant and storage of the pellet at −80°C.

### Radioligand binding assays

The affinities of reference standards and test compounds were determined via conventional competition and saturation radioligand binding assays. Competition assays were performed in round-bottom 96-well plates (Greiner) using standard binding buffer (50mM TrisHCl, 10mM MgCl_2_, 0.1mM EDTA, 0.1% fatty acid-free BSA, 1mM ascorbic acid, pH7.4) containing 3nM [^3^H]5-CT (44-158 Ci/mmol, PerkinElmer, Waltham, MA), serial dilutions of competitor (100μM to 0.01nM), and purified wild-type and mutant membranes. Pseudo-first order assumptions were met by using membranes at concentrations that bound <<10% of the radioligand added to each well. Non-specific binding was determined in the presence of 10 μM SB-699551. Plates were incubated in the dark for 2hr at RT and a PerkinElmer Filtermate harvester was used to collect membranes onto 0.3% PEI-treated GF/B glass fiber filtermats that were washed 4X with cold harvest buffer (50mM TrisHCl, pH7.4 at 4°C). The filters were dried, permeated with Meltilex scintillant (PerkinElmer, Waltham, MA), and counted on a Microbeta plate reader at 1min/well. Saturation binding assays were performed as above except that a serial dilution of [^3^H]5-CT (0.1nM to 25nM) was used. Bound cpm from competition experiments were analyzed in Prism (GraphPad Prism, San Diego, CA) using the Cheng-Prusoff correction to yield Ki equilibrium binding estimates. Equilibrium dissociation constants (K_D_) were fit directly from specific binding values using a one-site saturation model in Prism. Selectivity screens by the NIMH Psychoactive Drug Screening Program (PDSP) were performed as described (*5*). Validation assays for 5-HT_5A_R-Gi_1_ expression were performed as described above using 10nM [3H]-LSD (82.4 Ci/mmol, PerkinElmer, Waltham, MA).

### GTPγ[^35^S] loading assay

GTP*γ*[^35^S] assays were performed in 96-well plates in assay buffer (20 mM HEPES, 100 mM NaCl, 10 mM MgCl_2_, 1 mM EDTA, 1 mM DTT; pH 7.4) containing 10 μM GDP, 10μM GTP*γ*S (only for non-specific binding), 0.3 nM GTP*γ*[^35^S] (1250 Ci/mmol, PerkinElmer, Waltham, MA), and test or control ligands. In agonist mode, all ligands were screened at 32 μM. In antagonist mode, test ligands (100 μM) or control (10 μM SB-699551) were preincubated with receptor for 15-30 min at RT before an EC_80_ of 5-HT (1.0 μM) was added. GTP loading was initiated by addition of a premixture of cell membranes and WGA-SPA PVT beads (PerkinElmer, Waltham, MA) to a final concentration of 0.25mg beads/well and 15,000cpm 5-HT_5A_R-Gi_1_/well (as determined by [^3^H]-LSD binding). Plates were sealed and agitated at RT for 3hr (agonist mode) and 3-5hr (antagonist mode). Plates were counted in SPA mode in a PerkinElmer TriLux microbeta. Results (CPM) were normalized to 5-HT response in GraphPad Prism 5.0.

### Animals

Animal experiments were approved by the UCSF Institutional Animal Care and Use Committee and were conducted in accordance with the NIH Guide for the Care and Use of Laboratory animals. Adult (8-10 weeks old) male C56BL/6 mice were purchased from the Jackson Laboratory (strain #664). Mice were housed in cages on a standard 12:12 hour light/dark cycle with food and water ad libitum.

### Behavioral analyses

All ligands were dissolved in 20% ethanol at the desired concentration. For all behavioral tests, the experimenter was always blind to treatment. Animals were first habituated for 1 hour in Plexiglas cylinders and then tested 30 minutes after intrathecal (spinal cord CSF) injection of the compounds. Hindpaw mechanical thresholds were determined with von Frey filaments using the updown method (*42*). For the ambulatory (rotarod) test, mice were first trained on an accelerating rotating rod, 3 times for 5 min, before testing with any compound.

### Spared-nerve injury (SNI) model of neuropathic pain

Two of the three branches of the sciatic nerve were ligated and transected distally under isoflurane anesthesia, leaving the sural nerve intact. Behavior was tested 7 to 14 days after injury and *in situ* hybridization was performed one week post-injury.

### *In situ* hybridization

*In situ* hybridization was performed as previously described (*43*), using fresh DRG tissue from adult mice (8-10 week old) and following Advanced Cell Diagnostics’ protocol. All images were taken on an LSM 700 confocal microscope (Zeiss) and acquired with ZEN 2010 (Zeiss). Adjustment of brightness/contrast and introducing artificial colors (LUT) were done with Photoshop. The same imaging parameters and adjustments were used for all images within an experiment.

### Statistical analyses

All statistical analyses were performed with Prism (Graph Pad) using one-way ANOVA with Dunnett’s multiple comparison post-hoc test. Anatomical and behavioral data are reported as mean +/- SEM.

## RESULTS

### Homology model generation, vetting, and *in silico* screening

We used iterative modeling and testing to seek novel ligands selective for the 5-HT_5A_R (**Figure 1**). Because selective compounds of either class would be useful, we initially did not differentiate between agonists or antagonists. Moreover, as there is no crystal structure publicly available for the 5-HT_5A_R, we first built 1000 homology models based on the 5-HT_1B_R X-ray structure bound to ergotamine (PDB: 4IAQ; (*25*)) using Modeller (**Figure 1A**) (*27*). The 5-HT_1B_R was chosen because it shares 34% overall sequence identity with the 5-HT_5A_R and 49% sequence identity within the transmembrane regions. Ergotamine was retained in the modeling to ensure a ligand-competent orthosteric binding site.

The resulting models were evaluated for their ability to enrich 17 common lead-like 5-HTR agonists and antagonists (IUPHAR,(*44*)) over 1133 property matched decoys (**Figure 1B**). The best performing model by ligand enrichment was further optimized through energy minimization (**Figure 1C**) and selected for prospective virtual screening (**Figure 1D**). The best performing 5-HT_5A_R model was then screened against >6 million “lead-like” molecules (MW: 300-350, LogP: - 1 to 3.5) from the ZINC15 database (*37*). For many molecules, no successful pose was calculated; while 2,090,248 molecules were docked and scored. Top-ranked docked molecules were advanced if they were topologically dissimilar to ∼28,000 known aminergic ligands curated from the ChEMBL database (*38*) (ECFP4 Tc < 0.40). Finally, the top ranked 2,000 poses were visually inspected for unfavorable features that are sometimes missed by the docking scoring function, especially internal strain in the ligands and the occurrence of ligand hydrogen-bond donors that are not complemented by particular receptor acceptors, as described (*45*) (**Figure 1E**). Ultimately, 25 compounds, each representing a different chemotype (Table S1), were experimentally tested for binding to the 5-HT_5A_R (**Figure 1F**).

### Iterative docking and testing enables affinity maturation of a novel quinolone scaffold

The initial set of 25 docking hits and analogs were tested in binding assays of increasing stringency to probe their interactions with the orthosteric site. Testing at a single concentration of 100μM identified five chemotypes that displaced >50% of high affinity orthosteric agonist binding ([^3^H]5-CT) from the human 5-HT_5A_R (**Figure 2A-B**), corresponding to a 20% hit rate. In competition radioligand binding assays spanning eight orders of magnitude, the affinities of compounds 5A-6, 5A-7, 5A-9, 5A-15, and 5A-19 ranged from 12 to 42 μM (**Figure 2C, Table 1**) Although substantially weaker than the mid-nM antagonist SB-699551, several of these new ligands had antagonist-like activity in GTP-loading assays (**Figure 2D**). Based on their potency, target engagement in functional assays, and availability of analogs in the docking library, we sought to optimize the **5A-6** and **5A-9** scaffolds via a widely-used analog-by-catalog strategy (*8, 9, 47, 48*) **(Figure 2E-J**).

**Figure 2.**
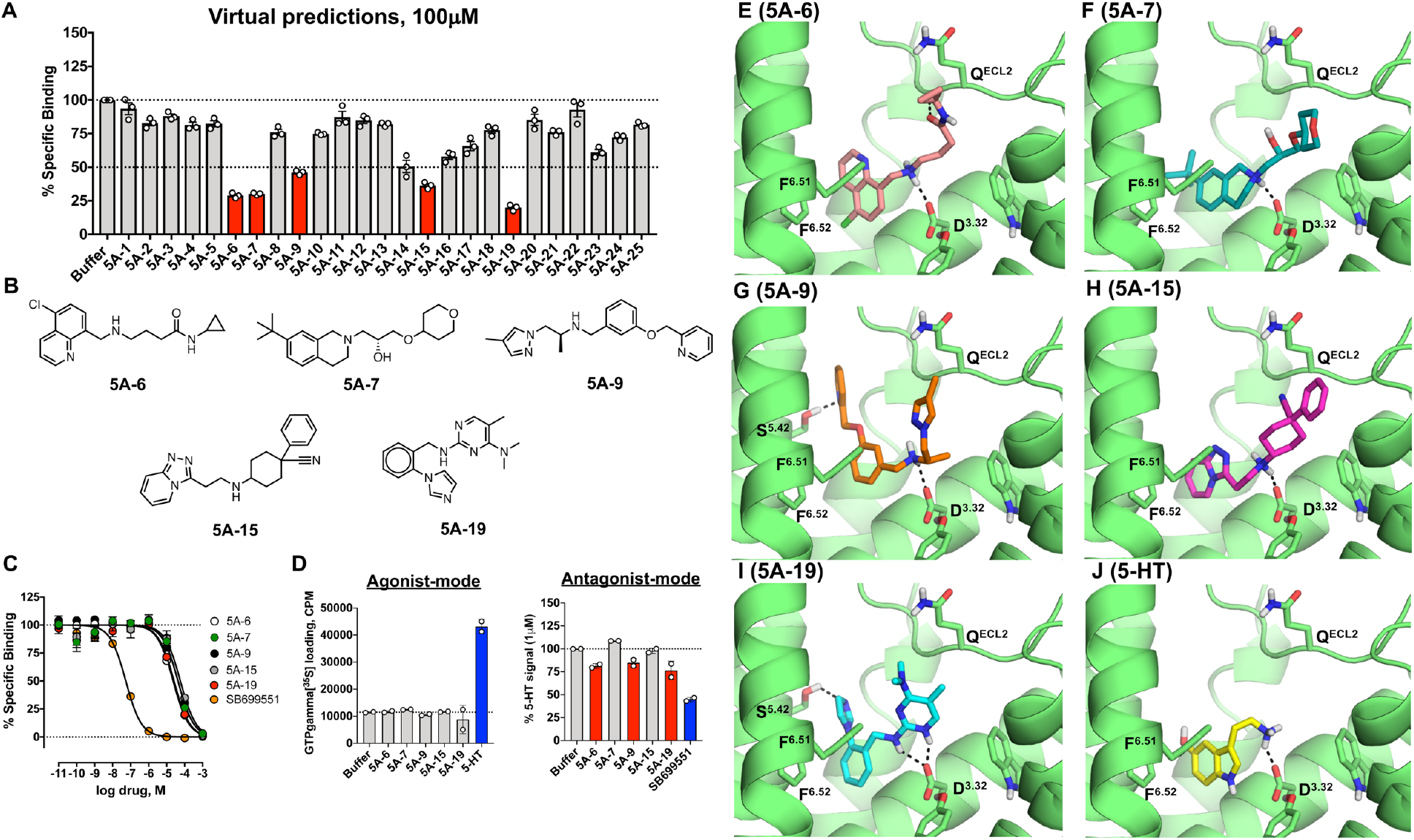
Library docking identifies novel 5-HT5AR chemotypes. **(A)** Single-point competition binding assay of 25 high-ranking docking molecules. At 100μM, 5 of the 25 (red bars) displaced >50% of the high affinity agonist [^3^H]5-CT. Data shown as mean ± SEM (n = 3, in experimental triplicate). **(B)** Chemical structures of the top five docking hits; each represents a different scaffold. **(C)** Concentration–response curves of [^3^H]5-CT displacement by the five docking hits, versus the known 5-HT5AR antagonist SB-699551. Data are mean ± SEM (n = 3-11, in experimental duplicate). **(D)** GTP*γ*[^35^S] loading assays for agonist (n=1 in duplicate) and antagonist (n=2 in triplicate) activity of five docking hits at 32 μM and 100 μM, respectively. Data shown are mean ± SD. Docked poses of compounds **5A-6 (E)**, **5A-7 (F)**, **5A-9 (G)**, **5A-15 (H)**, **5A-19 (I)** and 5-HT **(J)**. The 5-HT5AR is shown in green, and compounds are shown as capped sticks with colored carbons. Ballesteros-Weinstein numbering (*49*) is shown as superscript.

Docking of **5A-6** suggested that it interacts with conserved binding site residues including: a salt bridge between the ligand’s secondary amine and the D121^3.32^—a hallmark interaction between aminergic GPCRs and their ligands—and van der Waals contacts between the ligand’s halogenated quinoline and residues on TMs 3, 5 and 6, including C125^3.36^, A208^5.46^ and F302^6.52^ (**Figure 2E**). The docked pose also featured a new hydrogen bond interaction between the carbonyl group of the terminal ligand amide and the backbone of V194^ECL2^ (Extracellular Loop 2). Since the left hand side of 5A-6 coordinates many interactions common amongst biogenic amine receptors, and to all 5-HTRs in particular (*12*) (**Figure 2J**), we mainly explored substitutions on the right hand side of the molecule that extends towards the extracellular loops. The preliminary docking hit was optimized through several rounds of analoging within the ZINC15 database (*37*). In the first round, 4374 analogs were identified by a substructure similarity search using the core scaffold of **5A-6** against the “lead-like” subset of ZINC15. After removing all but the cationic molecules to preserve the D121^3.32^ interaction, these analogs were docked to the 5-HT_5A_R model. Analogs that maintained putatively key interactions observed for the parent molecule and that formed additional favorable contacts were selected for experimental testing. Similarly, docking of compound **5A-9** suggested that it hydrogen bonds with D121^3.32^ through its cationic nitrogen, as well as with S204^5.42^ (**Figure 2G**). A set of 2450 topologically similar analogs of **5A-9** were identified, as above, and docked to the 5-HT_5A_R model.

Together, 15 analogs of **5A-6** and 11 analogs of **5A-9** were prioritized for testing. Five analogs of 5A-6 outperformed the parent molecule in single-concentration testing at 32μM (**Figure 3A-B**; **Table 1**). Conversely, none of the **5A-9** analogs exhibited any improvement over the parent and this scaffold was not further pursued. Competition binding assays (**Figure 3C**) confirmed that **5A-6** analogs with a bulky rigidified ring system on the right-hand side, such as substituted furan (**5A6-8**) or piperidine (**5A6-1**) rings or a cyclic sulfone (**5A6-10**), bound with higher affinity than the parent molecule (**Figure 3B**). Compound **5A6-12** (1.5μM) showed the greatest improvement in affinity versus **5A-6** (12μM) (**Figure 3D**).

**Figure 3.**
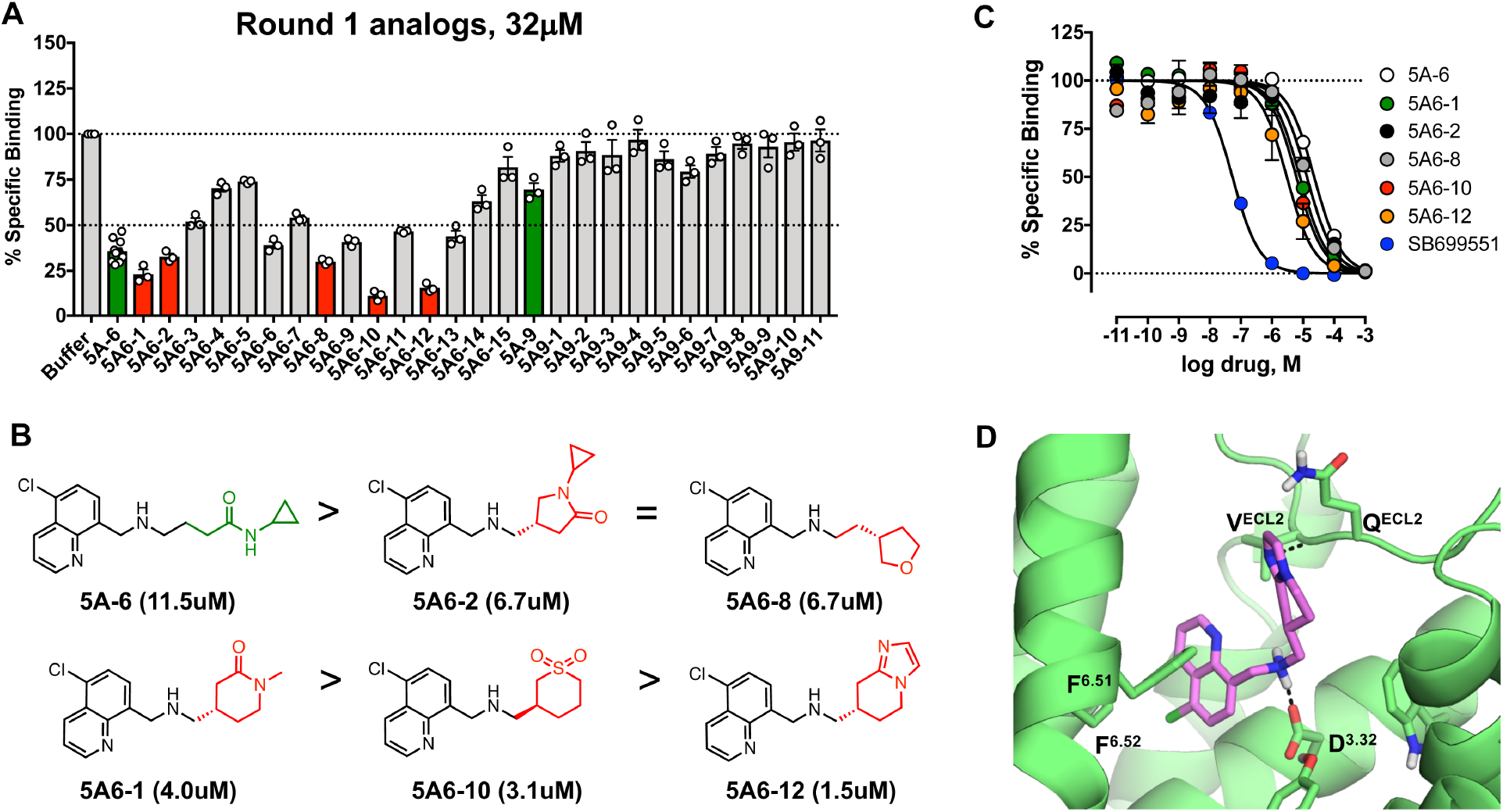
The 5A-6 series is identified as a candidate for optimization in first round of analoging. **(A)** Single-point competition binding assay of 15 analogs of compound **5A-6** and 11 analogs of compound **5A-9**. Five of the **5A-6** analogs (red bars) were capable of displacing binding of the high affinity agonist [^3^H]5-CT better than the parent compound **5A-6** (green bar). None of the **5A-9** analogs tested had better affinity than the parent compound (green bar). Each analog was tested at 32 μM. Data shown as mean ± SEM (n = 3, in experimental triplicate). **(B)** Chemical structures of the five active analogs relative to the parent compound **5A-6**. The variable group in each structure is colored in red. Ki values for each molecule are indicated below the structure. **(C)** Competition binding assays of the five active analogs relative to the parent compound **5A-6** and the known antagonist SB699551. Data shown as mean ± SEM (n=3-11, in experimental duplicate). **(D)** Docked pose of the analog **5A6-12**, which had the greatest improvement in affinity relative to **5A-6**. The 5-HT5AR is shown in green, **5A6-12** is shown as capped sticks with carbons colored pink. Ballesteros-Weinstein numbering is shown as superscript.

**Table 1:**
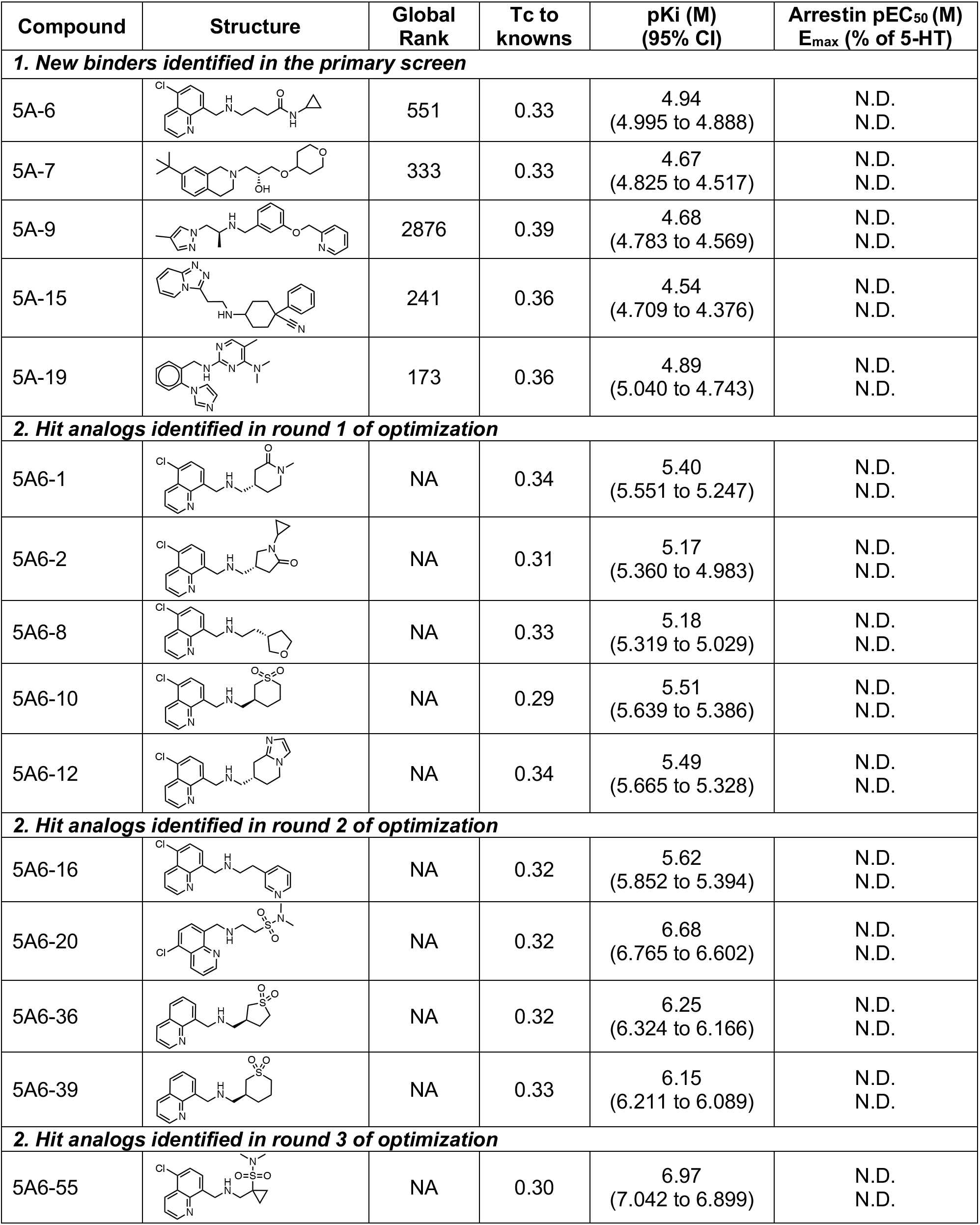
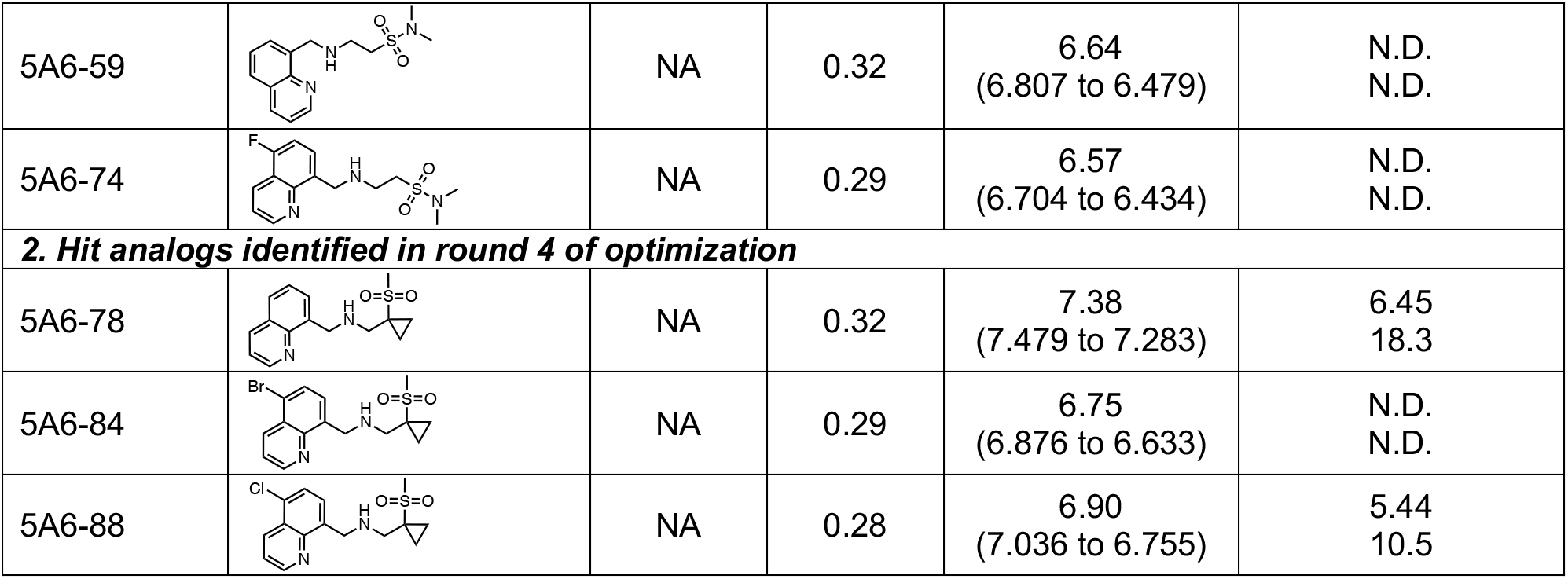
Optimization of compound affinity and potency for 5-HT_5A_R.

| Compound | Structure | Global Rank | Tc to knowns | pKi (M)<br>(95% CI) | Arrestin pEC<sub>50</sub> (M)<br>E<sub>max</sub> (% of 5-HT) |
| --- | --- | --- | --- | --- | --- |
| **1. New binders identified in the primary screen** |  |  |  |  |  |
| 5A-6 |  | 551 | 0.33 | 4.94<br>(4.995 to 4.888) | N.D.<br>N.D. |
| 5A-7 |  | 333 | 0.33 | 4.67<br>(4.825 to 4.517) | N.D.<br>N.D. |
| 5A-9 |  | 2876 | 0.39 | 4.68<br>(4.783 to 4.569) | N.D.<br>N.D. |
| 5A-15 |  | 241 | 0.36 | 4.54<br>(4.709 to 4.376) | N.D.<br>N.D. |
| 5A-19 |  | 173 | 0.36 | 4.89<br>(5.040 to 4.743) | N.D.<br>N.D. |
| **2. Hit analogs identified in round 1 of optimization** |  |  |  |  |  |
| 5A6-1 |  | NA | 0.34 | 5.40<br>(5.551 to 5.247) | N.D.<br>N.D. |
| 5A6-2 |  | NA | 0.31 | 5.17<br>(5.360 to 4.983) | N.D.<br>N.D. |
| 5A6-8 |  | NA | 0.33 | 5.18<br>(5.319 to 5.029) | N.D.<br>N.D. |
| 5A6-10 |  | NA | 0.29 | 5.51<br>(5.639 to 5.386) | N.D.<br>N.D. |
| 5A6-12 |  | NA | 0.34 | 5.49<br>(5.665 to 5.328) | N.D.<br>N.D. |
| **2. Hit analogs identified in round 2 of optimization** |  |  |  |  |  |
| 5A6-16 |  | NA | 0.32 | 5.62<br>(5.852 to 5.394) | N.D.<br>N.D. |
| 5A6-20 |  | NA | 0.32 | 6.68<br>(6.765 to 6.602) | N.D.<br>N.D. |
| 5A6-36 |  | NA | 0.32 | 6.25<br>(6.324 to 6.166) | N.D.<br>N.D. |
| 5A6-39 |  | NA | 0.33 | 6.15<br>(6.211 to 6.089) | N.D.<br>N.D. |
| **2. Hit analogs identified in round 3 of optimization** |  |  |  |  |  |
| 5A6-55 |  | NA | 0.30 | 6.97<br>(7.042 to 6.899) | N.D.<br>N.D. |
| 5A6-59 |  | NA | 0.32 | 6.64<br>(6.807 to 6.479) | N.D.<br>N.D. |
| 5A6-74 |  | NA | 0.29 | 6.57<br>(6.704 to 6.434) | N.D.<br>N.D. |
| **2. Hit analogs identified in round 4 of optimization** |  |  |  |  |  |
| 5A6-78 |  | NA | 0.32 | 7.38<br>(7.479 to 7.283) | 6.45<br>18.3 |
| 5A6-84 |  | NA | 0.29 | 6.75<br>(6.876 to 6.633) | N.D.<br>N.D. |
| 5A6-88 |  | NA | 0.28 | 6.90<br>(7.036 to 6.755) | 5.44<br>10.5 |

Beneficial chemical features identified in the first round of hit optimization, such as thiane– dioxide (**5A6-10**) and tetra-hydro-imidazo-pyridine (**5A6-12**) groups, were the basis for a second round of analoging. Similarity searches of the ZINC15 database followed by docking yielded a diverse set of 27 analogs (**Table S3**). Testing in competition radioligand binding assays under 32-fold more stringent conditions (1.0 μM) revealed four analogs with improved affinity (**Figure 4A-B**, **Table 1**). Binding affinity increased as much as 55-fold versus the parent molecule; analog rank order affinity (μM to nM): **5A6-16 > 5A6-39** > **5A6-36** > **5A6-20**, with the Ki of **5A6-20** reaching 208nM. As with Round 1, all top analogs of this set docked to hydrogen bond with the backbone amide of V194^ECL2^ (**Figure 4C**). As exemplified by **5A6-20**, affinity appeared to track with the nature of the sulfonyl variant - a sulfonamide substitution (**5A6-20**) was better than a cyclic sulfone (**5A6-36**/**39**; **Figure 4D**). Conversely, halogenation of the quinoline ring reduced affinity as exemplified by the close analogs **5A6-10** (3.1μM) and **5A6-39** (705nM).

**Figure 4.**
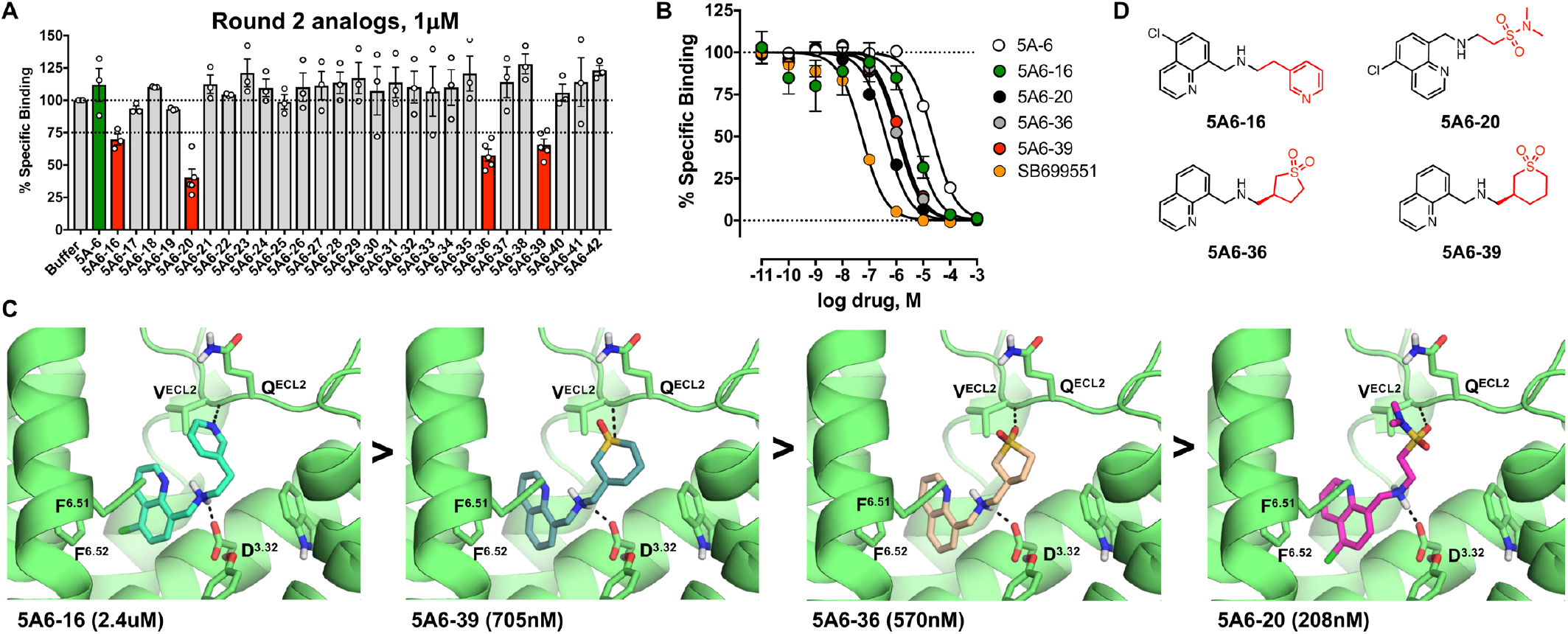
Optimization of chemical features identified in the first round leads to nanomolar ligands. **(A)** Single-point competition binding assay of 27 analogs. Four of the **5A-6** analogs (red bars) displaced the [^3^H]5-CT better than the parent compound **5A-6** (green bar). Each analog was tested at 1 μM. Data shown as mean ± SEM (n = 3-5, in experimental triplicate). **(B)** Competition binding assays of the four active analogs versus the parent **5A-6** and the known antagonist SB699551. Data shown as mean ± SEM (n = 3-11, in experimental duplicate). **(C)** Docked poses of the four active analogs (from left to right) **5A-16**, **5A6-39**, **5A6-36** and **5A6-20**. Ki values are indicated below the image. The 5-HT5AR is shown in green, and the molecules are shown as capped sticks with colored carbons. Ballesteros-Weinstein numbering is shown as superscript. **(D)** Chemical structures of the four active analogs. The variable group in each structure is colored in red.

### Further lead optimization and testing of the structural model

Cyclic sulfone and sulfonamide substituents conferred relatively high binding affinity to the quinoline ring and basic amine anchor scaffold, modeled to occur through hydrogen bonds with ECL2 residues (**Figure 4C**). In Round 3 we investigated changes to: i) the type and location of quinoline ring halogenation, including removing the halogen entirely (X); ii) the configuration of the sulfonamide (R2); iii) the configuration of the cyclic sulfone (R2); and iv) the type of hydrogen bond acceptor (R2) (**Figure 5A**, **Table S4**). We also explored the addition of hydrophobic substituents (R1). Analogs with a sulfonamide configuration similar to **5A6-20** exhibited the highest affinities (**Figure 5B-C**, **Table 1**). Compound **5A6-74** had an affinity similar to **5A6-59**, despite posing to interact with both D121^3.32^ and V194^ECL2^; while **5A6-59** is only modeled to hydrogen bond with D121^3.32^ alone. This may reflect the unfavorable effect of halogenation on **5A6-74**, an effect that is also apparent in comparing **5A6-10** and **5A6-39** (**Figures 3** and **4**). Compound **5A6-55**, which had the highest affinity of any analog tested thus far, resembled **5A6-20** but possessed an additional cyclopropyl group that is posed to make apolar contacts with W117^3.28^ and that re-orients the sulfonamide hydrogen bond between the mainchain nitrogen of Val^ECL2^ and a sulfonamide oxygen (**Figure 5C**). The advantages conferred by this cyclopropyl group may have overcome the negative effects of quinoline ring halogenation, which also occurs in **5A6-55**. Consistent with the modeled orientation, substitution of W117^3.28^ to alanine (W117A) decreased **5A6-55** binding affinity 10-fold, with only a 3.9-fold decrease on the endogenous ligand 5-HT that lacks hydrophobic substitution extending towards W117^3.28^ (**Figure 5D**). Intriguingly, the widely-used antagonist SB699551 bound threefold *better* to W117A^3.28^ than to the wild-type receptor (**Figure 5D**), supporting a different set of interactions and different overall pose within the binding pocket for the much larger SB699551 versus **5A6-55** and its congeners.

**Figure 5.**
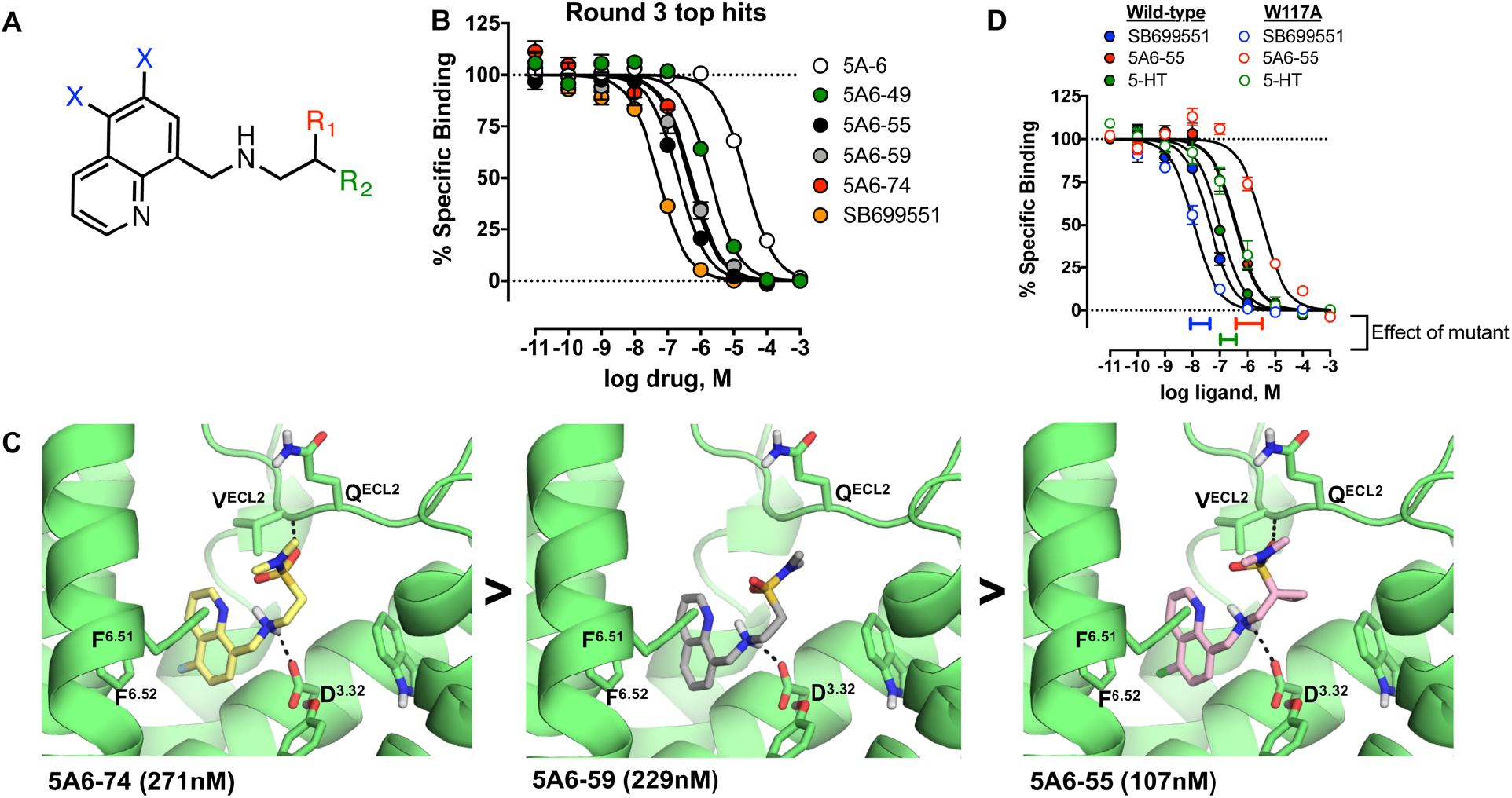
High affinity binding through sulfonamide substitutions and modeled engagement of ECL2. **(A)** Three regions (X, R1, R2) targeted in the third round of optimization are highlighted on the structure of the anchor scaffold. **(B)** Competition binding assays of the four active analogs identified in this round versus the parent **5A-6** and the known antagonist SB699551. Data shown as mean ± SEM (n=3-11, in experimental duplicate). **(C)** Docked poses of the three most potent analogs (from left to right) **5A6-74**, **5A6-59**, and **5A6-55**. Ki values are indicated below the image. The 5-HT5AR is shown in green, and the molecules are shown as capped sticks with colored carbons. Ballesteros-Weinstein numbering is shown as superscript. **(D)** Competition binding assays of **5A6-55**, the known antagonist SB-699551, and the endogenous ligand 5-HT at Wild-type 5-HT5AR and the single-point mutant W117^3.28^A. Data shown as mean ± SEM (n=3, in experimental duplicate).

A final round of analogs combined minor changes to the halogenation state of the quinoline with modifications to the sulfone group, to the size and position of the hydrophobic ring, and to the hydrogen bond acceptor (**Table S5**). Converting the sulfonamide to a methyl-sulfone, retaining the cyclopropyl ring, while dehalogenating the quinoline ring yielded the 41nM **5A6-78** (**Figure 6**, **Table 1**). Consistent with trends seen throughout the series, bromination (**5A6-84**) and chlorination (**5A6-88**) of the quinoline ring of **5A6-78** at the 6-position decreased affinity 4.2- and 3.1-fold, respectively (**Figure 6B**). From the docking poses, these decreases may reflect the loss of a hydrophobic contact with A208^5.46^ on TM5. As with the highly similar analog **5A6-55**, but larger in its overall effect, mutating W117^3.28^ to alanine decreased **5A6-78** binding affinity more than 25-fold (**Figure 6C**), supporting both an important interaction with this side chain and the docking pose. While W117^3.28^ is fairly conserved amongst 5-HTR subtypes, it is substituted in 5-HT_1A_R (Phe), 5-HT_4_R (Arg), and 5-HT_7_R (Phe).

**Figure 6.**
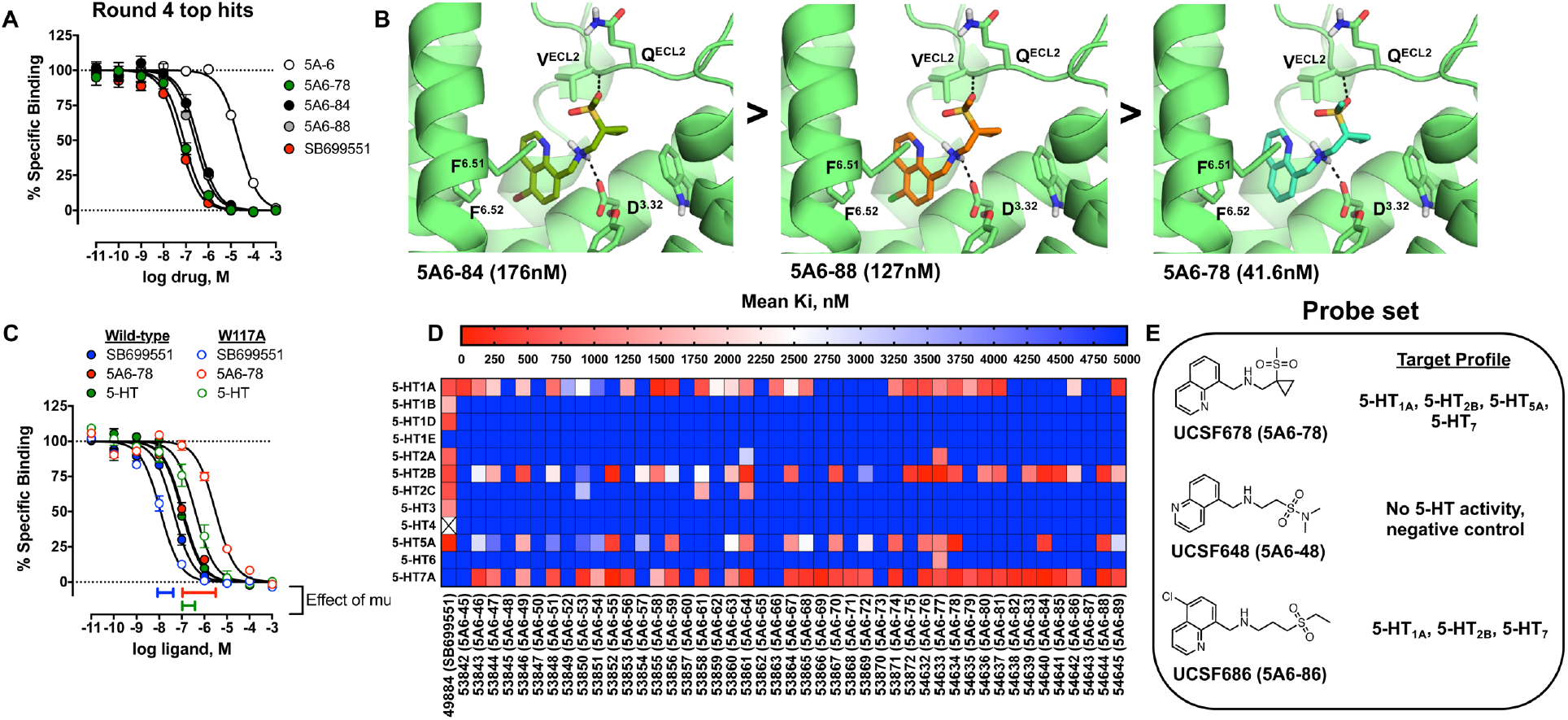
High affinity analogs have a more restricted binding profile across the 5-HT receptor family. **(A)** Competition binding assays of the four most active analogs identified in round 4 of analoging relative to the parent compound 5A-6 and the known antagonist SB699551. Data shown as mean ± SEM (n=3-11, in experimental duplicate). **(B)** Docked poses of the three most potent analogs (from left to right) **5A6-84**, **5A6-88**, and **5A6-78**. Ki values are indicated below the image. The 5-HT5AR is shown in green, and the molecules are shown as capped sticks with colored carbons. Ballesteros-Weinstein numbering is shown as superscript. **(C)** Competition binding assays of **5A6-78**, the known antagonist SB-699551 and the endogenous ligand 5-HT at Wild-type and the single-point mutant W117^3.28^A. Data shown as mean ± SEM (n=3-6, in experimental duplicate). **(D)** Affinity panel with Ki values (nM) of analogs from rounds 3 and 4 against all members of the 5-HT receptor family. **(E)** Target profile of **UCSF678** probe set.

### Comprehensive affinity profiling reveals that the novel quinoline/sulfone scaffold confers a more restricted binding profile than SB-699551

In collaboration with the NIMH Psychoactive Drug Screening Program (PDSP, https://pdspdb.unc.edu/pdspWeb), we comprehensively profiled round 3 and round 4 analogs across twelve 5-HTRs, assessing selectivity versus the widely-used reagent, SB-699551. SB-699551 exhibited appreciable affinity (Ki<2μM) for many 5-HTR subtypes including 5-HT_1A_R, 5-HT_1B_R, 5-HT_1D_R, 5-HT_2A_R, 5-HT_2B_R, 5-HT_2C_R, and 5-HT_3_ (ion channel), in addition to the 5-HT_5A_R (**Figure 6D)**. Conversely, most of our analogs lacked affinity for 5-HT_1B_R, 5-HT_1D_R, 5-HT_2A_R, 5-HT_2C_R, and 5-HT_3_. Unique to our analog series was a gain in affinity for 5-HT_7A_, which is predicted to have a nearly identical orthosteric binding pocket to 5-HT_5A_R. Our highest affinity compound, **5A6-78**, exhibited a marked improvement over SB-699551, with binding restricted to just 5-HT_1A_R, 5-HT_2B_R, and 5-HT_7A_R, in addition to 5-HT_5A_R (**Figure 6D**). We will refer to **5A6-78** as **UCSF678** from here on.

To control for the remaining off-target activity of **UCSF678**, we sought molecules that closely resembled **UCSF678** structurally but lacked binding to 5-HT_5A_R or other off-targets. Such molecules, used in counterpoint to **UCSF678**, can act as “probe pairs” that disentangle the on-from off-target effects of the probe. Two close analogs emerged: **5A6-48** (hereafter referred to as **UCSF648)**, which has no measurable effect on any of the 5-HTR receptor subtypes, and **5A6-86** (hereafter referred to as **UCSF686)**, which lost affinity at 5-HT_5A_R (>10,000nM) but not at 5-HT_1A_R, 5-HT_2B_R and 5-HT_7_R (**Figure 6E**). Used together in seeking phenotypic effects in cells, tissues, or organisms, the “probe triple” of **UCSF678, UCSF648**, and **UCSF686** controls for general, non-5-HTR binding of the series (**UCSF648**) and for the off-target engagement of 5-HT_1A_R, 5-HT_2B_R and 5-HT_7A_R (**UCSF686**). A comprehensive off-target analysis for these probes can be found in Supplementary Figure 1.

**Weak partial agonism and arrestin bias of UCSF678, and GPCRome-wide activity profiling.** Arrestin recruitment is a sensitive screen for GPCR activity especially when studying orphan and understudied GPCRs (*4*). Here we tested the initial 25 docking hits and subsequent **5A-6** series analogs for their ability to recruit *β*-arrestin 2 relative to full agonists. Tested at 10 μM (**Figure 7A**) and confirmed as concentration-response curves (**Figure 7B**), the analogs **5A6-36**, **5A6-39**, and **UCSF678** were partial agonists for *β*-arrestin 2 recruitment. **UCSF678** had the most robust signal, recruiting *β*-arrestin 2 to 17% of 5-HT and 5-CT controls. Counter-screens at the Gi/o family transducer G*α*oA failed to detect activation relative to the 5-HT control (**Figure 7C and** **D**), suggesting that **UCSF678** is *β*-arrestin-biased.

**Figure 7.**
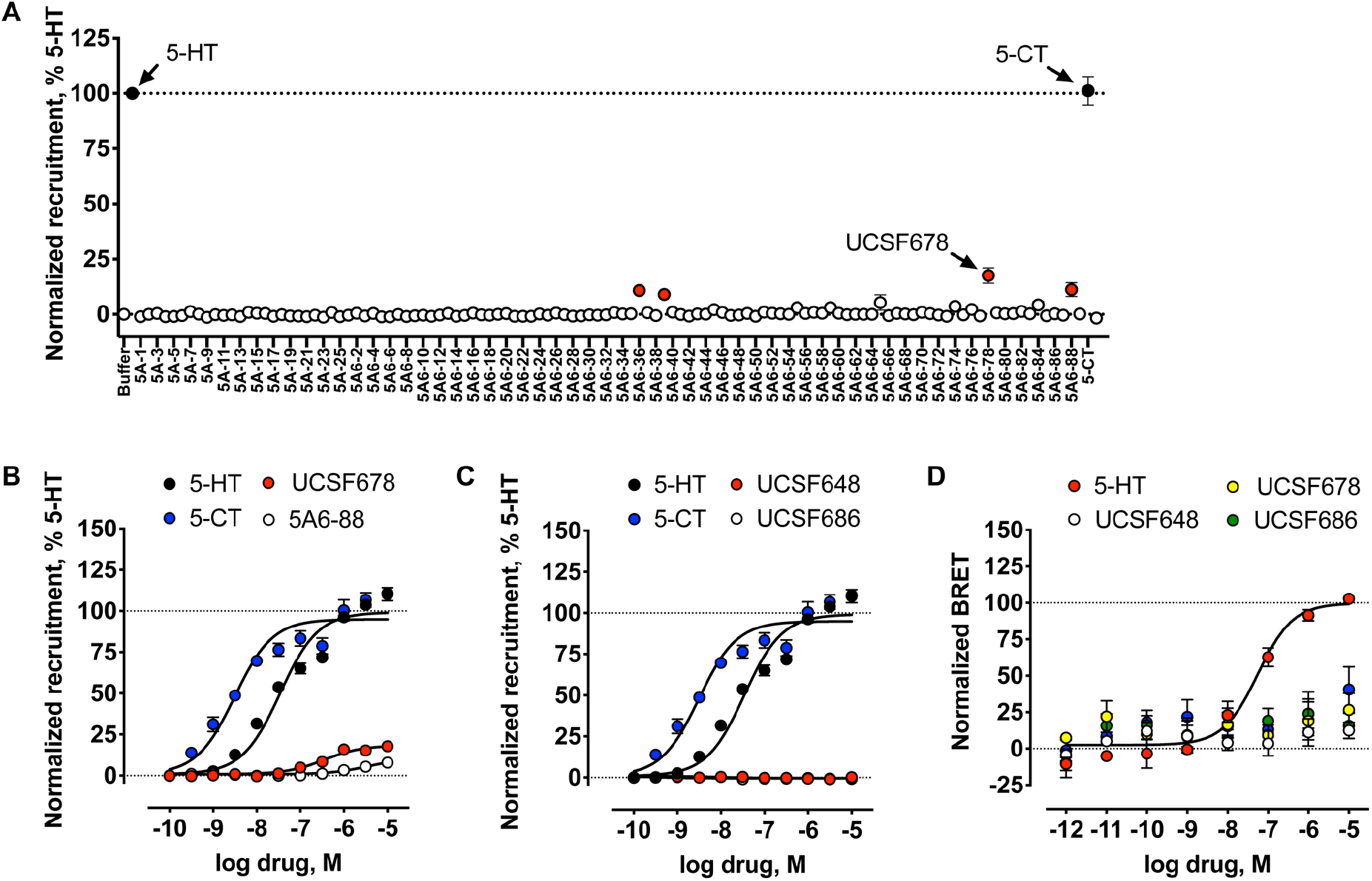
Comprehensive activity profiling reveals that select analogs are arrestin-biased partial agonists. **(A)** Tango beta-arrestin 2 recruitment assays (*4*) testing the activity of initial virtual docking hits and subsequent **5A-6** series analogs at 10μM at the human 5-HT5AR. Values are normalized to the full agonist serotonin (5-HT, 100%) and presented as mean ± s.e.m. of three to five experiments in quadruplicate. **(B,C)** The activities of probe molecules **5A6-48**, **5A6-78**, and **5A6-86** for beta-arrestin 2 recruitment were confirmed in concentration-response curves. Emax values of 5-HT for each plate were calculated via three-parameter logistic fits in GraphPad Prism and used to normalize the raw luminescence values of test ligands. Baseline luminescence values were shared between concentration response curves on the same 384-well plate. Normalized mean luciferase values (%) were combined across experiments and re-fit to the three-parameter model sharing baseline values. Data shown are mean ± s.e.m. of four to eleven experiments in quadruplicate. **(D)** Testing activation of the G*α*oA pathway by probe molecules **UCSF648**, **UCSF678**, and **UCSF686**, and **5A6-88** and the reference full agonist 5-HT. Emax values of 5-HT for each plate were calculated via three-parameter logistic fits in GraphPad Prism and used to normalize the BRET2 ratios of test ligands. Data shown as mean ± SEM (n=4, in experimental duplicate).

For completeness, we screened the probe set molecules **UCSF648**, **UCSF678**, and **UCSF686** across >300 GPCRs in the Tango assay to detect activation across the GPCRome (*4*). We followed-up several GPCRome hits in concentration-response curves to reveal that our probe set was largely inactive. Specifically, **UCSF648** weakly activated ADRA2A and MTNR1A (**Supplementary Figure 2A-C**), UCSF686 weakly activated CXCR7 (**Supplementary Figure 2D,E**), and the probe molecule **UCSF678** activated the D2 dopamine receptor and 5-HT_2C_R (INI isoform) with low potencies (>1.0 uM, **Supplementary Figure 2F-H**).

### The widely used antagonist SB-699551 has liabilities as a chemical probe

SB-699551 is a tool molecule that, over the last eight years, has been widely used to investigate the *in vivo* and cellular roles of the 5-HT_5A_ receptor (*16-18, 20, 21, 24, 50-57*). Use in these studies is predicated not only on its relative selectivity—now in doubt (**Figure 6D**)—but also on its behavior in assays. Compared to **UCSF678,** SB-699551 exhibited near complete concentration-dependent inhibition of luminescence, as early as 20min after drug addition, in the absence of stimulation of the 5HT_5A_ receptor, suggesting that SB-699551 is an inhibitor of the luminescence assay itself (**Supplementary Figure 3**). Moreover, brightfield illumination of HEK293T cells treated overnight with high concentrations of SB-699551 suggested extensive cytotoxicity compared to the vehicle control and to **UCSF678**. Since many GPCR assays use luminescence as a proxy for activity (e.g. Tango *β*-arrestin 2 recruitment, BRET2 G protein activation, and GloSensor cAMP), SB-699551 may show antagonist activity against many GPCRs, at least at these concentrations. This is an artifact that has been seen previously in other contexts (*58, 59*). These results suggest caution in interpreting the activity of this compound in pharmacological studies.

### Antinociceptive behavior of new chemical probes

5-HT_5A_R antagonism has been associated with nociception and/or mechanical hypersensitivity/allodynia in mice (*51, 55, 57*); however, the off-target profiles of previously used antagonists confound ready interpretation. With a probe set that controls for activity at different 5-HTR receptors (**Figure 6D**), we sought to interrogate the contributions of 5-HT_5A_R and other 5-HTR subtypes to nociception. Here we tested the analgesic properties of our 5A-6 probe set in the context of neuropathic pain via the spared nerve injury (SNI) model, in which 2 out of 3 branches of the sciatic nerve are cut (*60*). To selectively investigate the contribution of spinal cord 5-HT_5A_R, we administered ligands intrathecally. Unlike the *inactive* control probe **UCSF648**, which lacked effect in the SNI mice (**Figure 8A**), intrathecal injection of **UCSF678** or the highly related control probe **UCSF686**, which is devoid of 5-HT_5A_R activity, substantially increased the mechanical thresholds ipsilateral to the injury side versus vehicle (**Figure 8**). Surprised by these findings, we expanded the study to other probes in our set to verify that 5-HT_5A_R affinity is indeed nonessential. **5A6-56** exhibited anti-allodynic effects comparable to **UCSF-678** despite having a binding profile restricted to just the 5-HT_1A_R and 5-HT_7A_R (one-way ANOVA; p<0.0001). Conversely, **5A6-88,** which shares the binding profile of **UCSF678** but lacks 5-HT_1A_R affinity, was not anti-allodynic, consistent with the primacy of the 5-HT_1A_R in rodent models of pain (*61–64*). Interestingly, intrathecal **5A6-56**, **UCSF678** and **UCSF686** had no visible effect on baseline mechanical thresholds, i.e. in the absence of nerve injury (**Figure 8B**). Importantly, intrathecal administration of **5A6-56**, **UCSF678** and **UCSF686** was devoid of sedating effects on the rotarod test (**Figure 8C**), indicating that their anti-allodynic effects were not the result of motor impairment. Taken together, 5-HT_5A_R modulation is non-sufficient in a rodent neuropathic pain model when we disentangle the on-from off-target effects of **UCSF678**, highlighting the importance of property-matched control probes.

**Figure 8:**
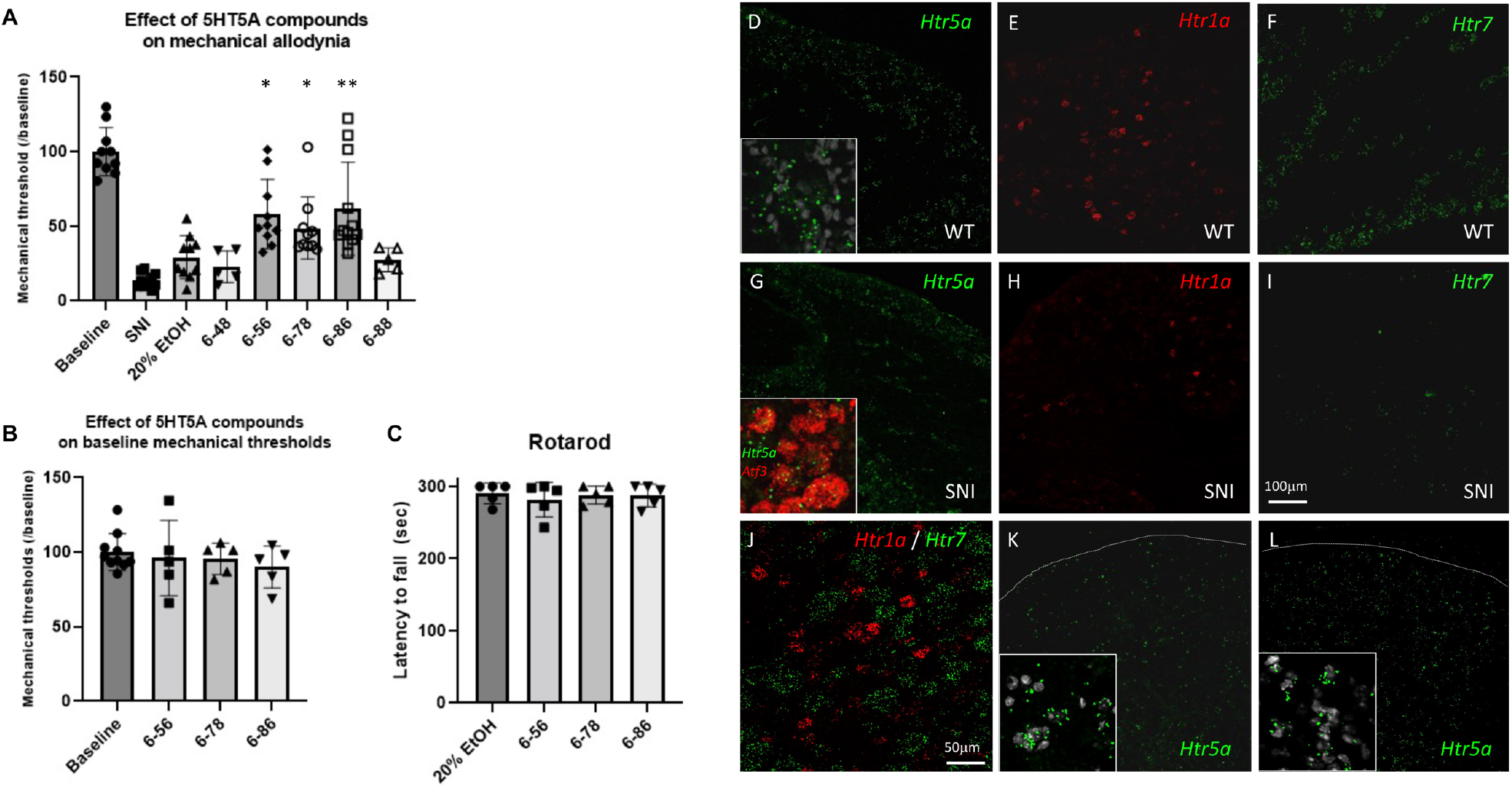
Intrathecal injection of the 5A-6 probe set is anti-allodynic. **(A)** The **5A6-56**, **UCSF678** and **UCSF686** ligands reduced the mechanical hypersensitivity that develops following sciatic nerve injury (SNI), compared to vehicle (20% ethanol). In contrast, the **UCSF648** and **UCSF688** ligands were ineffective. Data shown are mean ± SEM; One way ANOVA with Dunnett’s multiple comparison post-hoc test was performed to compare the effect of the various ligands to the vehicle control (20% ethanol); * p<0.05; **p<0.01. **(B-C)** None of the ligands tested altered the baseline mechanical thresholds in absence of injury (B) or the motor performance in the rotarod test (C). **(D-L)** *In situ* hybridization illustrates mRNA coding for the 5-HT receptor subtypes 5A (green; D, G), 1A (red, E, H and J) and 7 (green, F, I and J). These receptors are expressed at various levels in sensory neurons before (D-F) or after (G-I) partial sciatic nerve injury (SNI). Note that the 1A and 7 subtypes are expressed in non-overlapping subsets of DRG neurons (J). The 1A and 7 (but not 5A) subtypes are downregulated in DRG after SNI. **(K-L)** The 5-HT_5A_R mRNA is also expressed in the dorsal horn of the spinal cord, both before (K) and after (L) peripheral nerve injury. Scale bar is 100 um for all panels except G, where it is 50 um.

We used *in situ* hybridization to confirm that the 5-HT receptor targets of our 5A-6 probe set were expressed at the level of the spinal cord as well as the dorsal root ganglia (DRG), where the cell bodies of the sensory neurons that transmit the “pain” message to the spinal cord reside. Consistent with previous studies (*19, 20, 65*), we found that the 5-HT_5A_ subtype is expressed in a wide variety of spinal cord and DRG neurons (**Figure 8**). Interestingly, the 5-HT_1A_ and 5-HT_7_ subtypes were also expressed in DRG neurons but in non-overlapping subsets (**Figure 8**). Furthermore, we found that the expression level of 5-HT_7_R decreased dramatically in DRG neurons seven days after SNI (**Figure 8I**), whereas 5-HT_5A_R and 5-HT_1A_R remained unchanged (**Figure 8G-H**). Somewhat surprisingly, we could not detect the 5-HT_2B_ receptor subtype in DRG neurons, before or after SNI. Taken together, we conclude that intrathecal administration of our novel 5A-6 probe set can reduce the mechanical allodynia that develops following peripheral nerve injury, likely via an action at sites that include both the spinal cord and the primary afferent neurons (*66, 67*).

## DISCUSSION

Here we generated a novel series of 112 compounds from five rounds of integrated docking against a homology model, analoging, and empirical testing. From this approach emerged a 41nM subtype-selective chemical probe (**UCSF678**) with weak partial agonism and beta-arrestin-bias against the 5-HT_5A_R, along with two close analogs, **UCSF648** and **UCSF686,** that together control for off-target effects. Consistent with hit rates for previous GPCR virtual screening campaigns (*7*), 20% of the docking predictions were confirmed experimentally, a hit-rate that was maintained during increasingly stringent affinity maturation to select for potent and selective compounds.

A goal of the study was to find chemical probes with enhanced selectivity versus the widely used 5-HT_5A_R antagonist SB-699551. We did not initially screen for selectivity across 5-HTRs; instead, we sought chemically novel scaffolds exemplified by compound **5A-6**, enriching for those that exploited different interactions within the orthosteric pocket. Precedence for this comes from reports that unrelated classes of ligands can bind to the same receptor binding pocket (*47, 68–71*) and from library docking campaigns where chemical novelty translates into both sub-type and functional selectivity (*48, 72*). Thus, by advancing a novel quinoline/sulfone scaffold we hoped to exploit a different set of binding pocket residues that varied in distribution across 5-HTR subtypes. Compared to promiscuous 5-HTR ligands such as serotonin itself, the docked poses of our highest affinity compounds like **UCSF678** and **5A6-88** extend into upper regions of the binding pocket. We exploited such modeled interactions to further reduce their off-target activity, while also exploiting interactions with a conserved residue (W117^3.28^) for high affinity binding of **UCSF678** and other analogs (via cyclopropane). While mutagenesis experiments suggest that interactions with this tryptophan are important for affinity, we do note that this residue is conserved at all subtypes except 5-HT_1A_R (Phe), 5-HT_4_R (Arg,) and 5-HT_7_R (Phe), likely contributing to the off-target binding of our series. Despite this limitation, the chemical novelty of the scaffolds explored led to substantially improved selectivity (**Figure 6D**).

Using a combination of transcriptomics, mouse genetics, and small molecules, previous studies have begun to illuminate the *in vitro* pharmacology and *in vivo* roles of the 5-HT_5A_R, revealing its potential for clinical targeting in CNS diseases and pain (see (*13, 15*) for review). However, many of these studies used SB-699551 as a “selective” 5-HT_5A_R antagonist. This is understandable as it was a readily accessible, best-in-class molecule. Here we find that SB-699551 has liabilities as a chemical probe: it has substantial affinity for many 5-HTRs (**Figure 6D**), it lacks inactive property-matched controls, it artifactually decreases assay luminescence, and it appears cytotoxic at relevant concentrations. This can have important repercussions both *in vitro* and *in vivo*. For instance, the Kassai *et al*. study revealed that SB-699551 caused sedation that confounded interpretation of its anxiolytic effect. Two other antagonists, ASP5736 and A-843277, are reported to be selective enough for assigning 5-HT_5A_R function in animal models (*24, 73, 74*). However, neither antagonist is readily accessible, and neither is controlled by a “probe pair” for inevitable off-target activities. Meanwhile, **UCSF678** has i) an affinity for 5-HT_5A_R resembling that of the previous molecules (42nM, **Figure 6A**), ii) a more restricted off-target profile across 5-HTRs (**Figure 6D and 7**) and >300 GPCRs (**Figure 8**), iii) no cytotoxicity or assay interference (Figure 9), and iv) “probe pair” molecules with which to control for its off-target activities (**Figure 6E**). Accordingly, we are making the “probe triple” of **UCSF678**, **UCSF648**, and **UCSF686** readily available to the community, via the Sigma probe collection (registry numbers numbers in progress)

Our *in vivo* studies highlight the usefulness of property-matched probe pair molecules for disentangling the on-from off-target effects of chemical probes to correctly assign biological functions to understudied GPCRs. Specifically, our use of **UCSF678** analogs that bind different off-targets previously associated with analgesia (e.g. 5-HT_1A_R and 5-HT_7A_R (*75–78*)) call into question the extent to which 5-HT_5A_R signaling is essential for rodent nociception and/or analgesia. If such experiments were conducted without the full set of control probes used here, a very different conclusion may have been reached. Echoing the arguments of others (*79, 80*), we suggest that wherever possible chemical probe sets should be extended to include close analogs that lack activity at the intended target but retain off-target activities of the lead probe. Given that SB-699551 is far more promiscuous and poorly controlled relative to the probes developed here (**Figure 6D**), previous functions assigned to the 5-HT_5A_R merit reconsideration.

## CONCLUSION

The 5-HT_5A_R remains the least understood serotonin receptor, despite its association with multiple CNS disorders and in nociception. Indeed, it could be due to our lack of good pharmacological tools that the receptor has been implicated in quite so many disorders, from motor coordination and control, to exploratory behavior and anxiety, neuroendocrine function, learning and memory, emotion, brain development, the psychotropic effects of LSD, sleep, circadian rhythm, bladder function, arterial chemoreception (*13*),(*15*), memory stabilization (*16*) and amnesia (*17*), and, investigated here, nociception (*51, 55, 57*) and antiallodynia (*20, 21*). Here we used iterative rounds of structure-guided docking, pharmacological testing, and optimization to discover a new chemical probe for the 5-HT_5A_R, **UCSF678**, that is substantially more selective than existing 5-HT_5A_R antagonists, and better behaved in cell culture and *in vitro* than existing reagents. We combined this probe with two close analogs that are inactive on the 5-HT_5A_R, but control for activity on other 5-HTR subtypes, and for other off-targets to which the more general chemotype might bind. Whereas **UCSF678** is active against pain in a mouse model, the activities of the probe triple and of other close analogs—each with different profiles against the 5-HT receptor subtypes—suggest that the 5-HT_5A_R does not have a major role in treating pain, but rather this flows from other serotonergic receptors, likely the 5-HT_1A_R. As with other chemical probes for understudied GPCRs (*8, 9*), these molecules will help to more accurately assign biological functions to these targets. More broadly, the close cycle of modeling, docking, and *in vitro* pharmacology used here may find broad utility in the field. To that end, we make the docking libraries on which we drew (http://zinc15.docking.org and http://zinc.20.docking.org), the cell constructs and assays used for the *in vitro* pharmacology, and the 5-HT_5A_R probe triple (registry numbers in progress with Sigma-Millipore) openly available to the community.

## Acknowledgements

Supported by US NIH grants U24DK1169195 (to B.L.R & B.K.S), R01 MH112205, and the NIMH Psychoactive Drug Screening Contract (to B.L.R.); DARPA HR0011-19-2-0020 (to B.K.S., B.L.R. and A.I.B) R35 GM122481 (to B.K.S.); R35 NS097306 and Open Philanthropy (to A.I.B.); and K01 MH109943 (to R.T.S).

## Supporting Information for

**Supplementary Figure 1.**
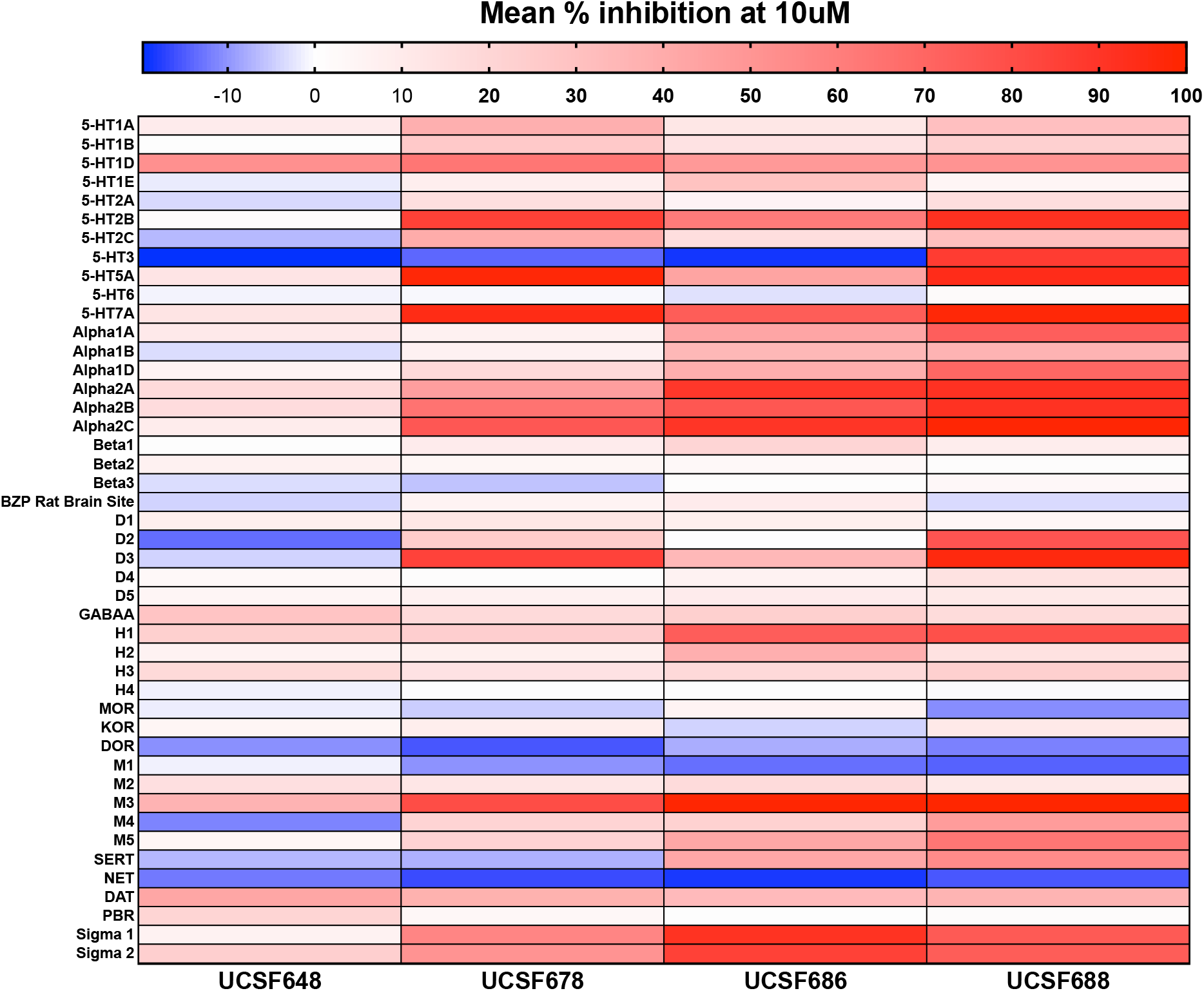
Expanded off-target engagement screen for 5A6-48, 5A6-78, and 5A6-86 probe set molecules across human and rodent receptors, channels, and transporters. Selectivity screens by the NIMH Psychoactive Drug Screening Program (PDSP) were performed as previously described (*5*). Heat map shows mean percent inhibition values performed in quadruplicate at 10μM.

**Supplementary Figure 2.**
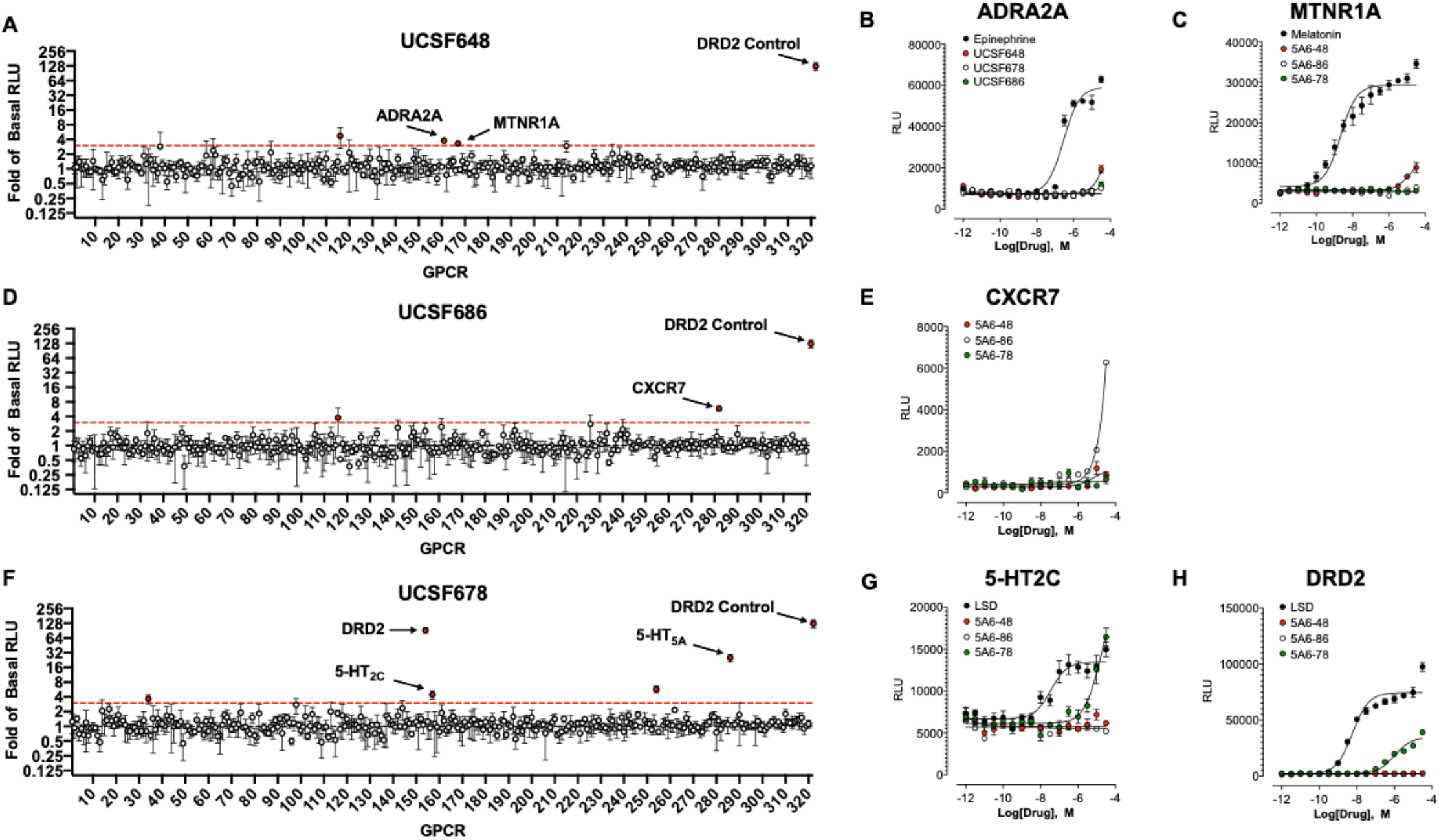
Comprehensive GPCRome profiling reveals limited off-target activity of probe set molecules. Tango screens of >300 GPCRs were conducted to assess off-target activity of probe molecules (A) 5A6-48, (B) 5A6-78, and (C) 5A6-86. All compounds were screened at 10μM, with the dopamine D2 receptor serving as the positive control. Confirmation of primary screen hits shown as concentration-response curves (right). Data shown as mean ± SEM (n=1, in experimental quadruplicate).

**Supplementary Figure 3.**
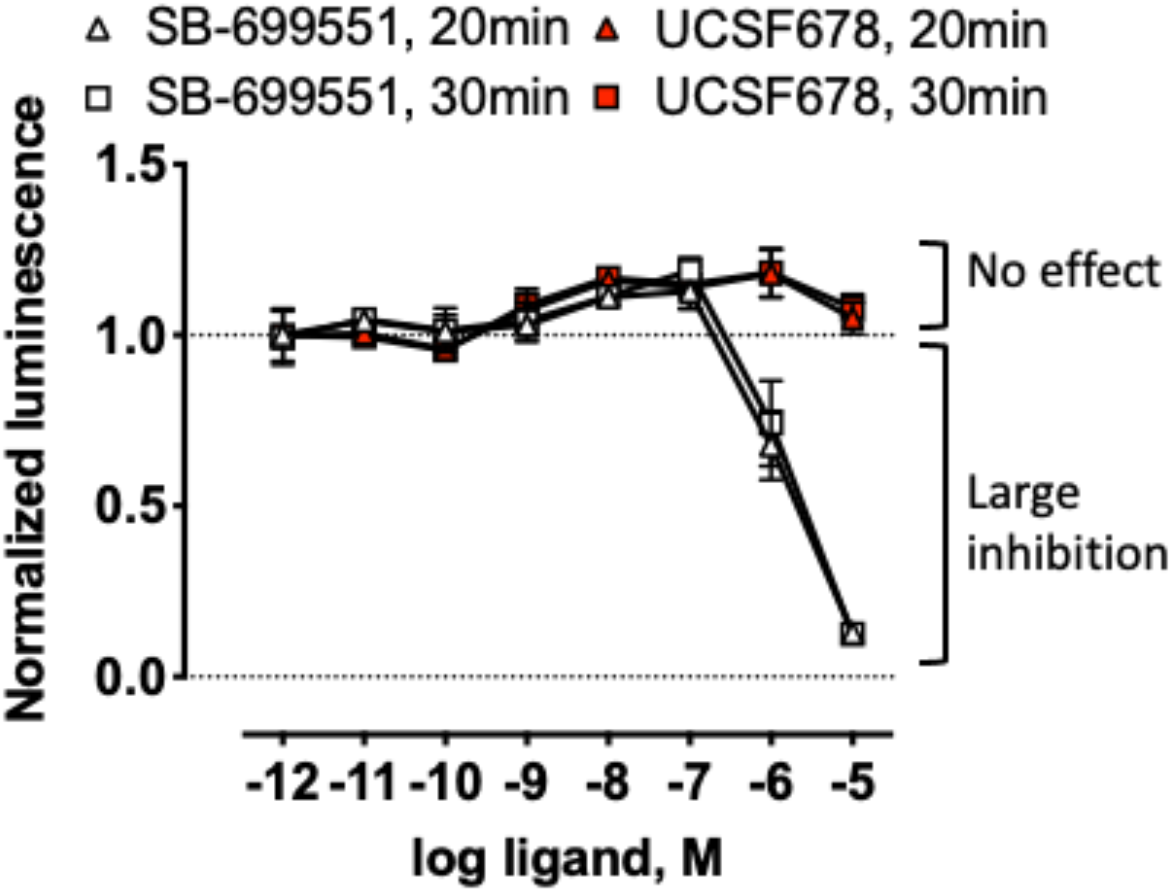
The commercially available antagonist SB-699551 decreases luminescence in cellular assays unstimulated by agonist. Shown are normalized concentration-response curves for RLuc8-based luminescence in the presence of 5A6-78 and SB-699551. Data represent mean ± SEM (n=2).

**Table S1:**
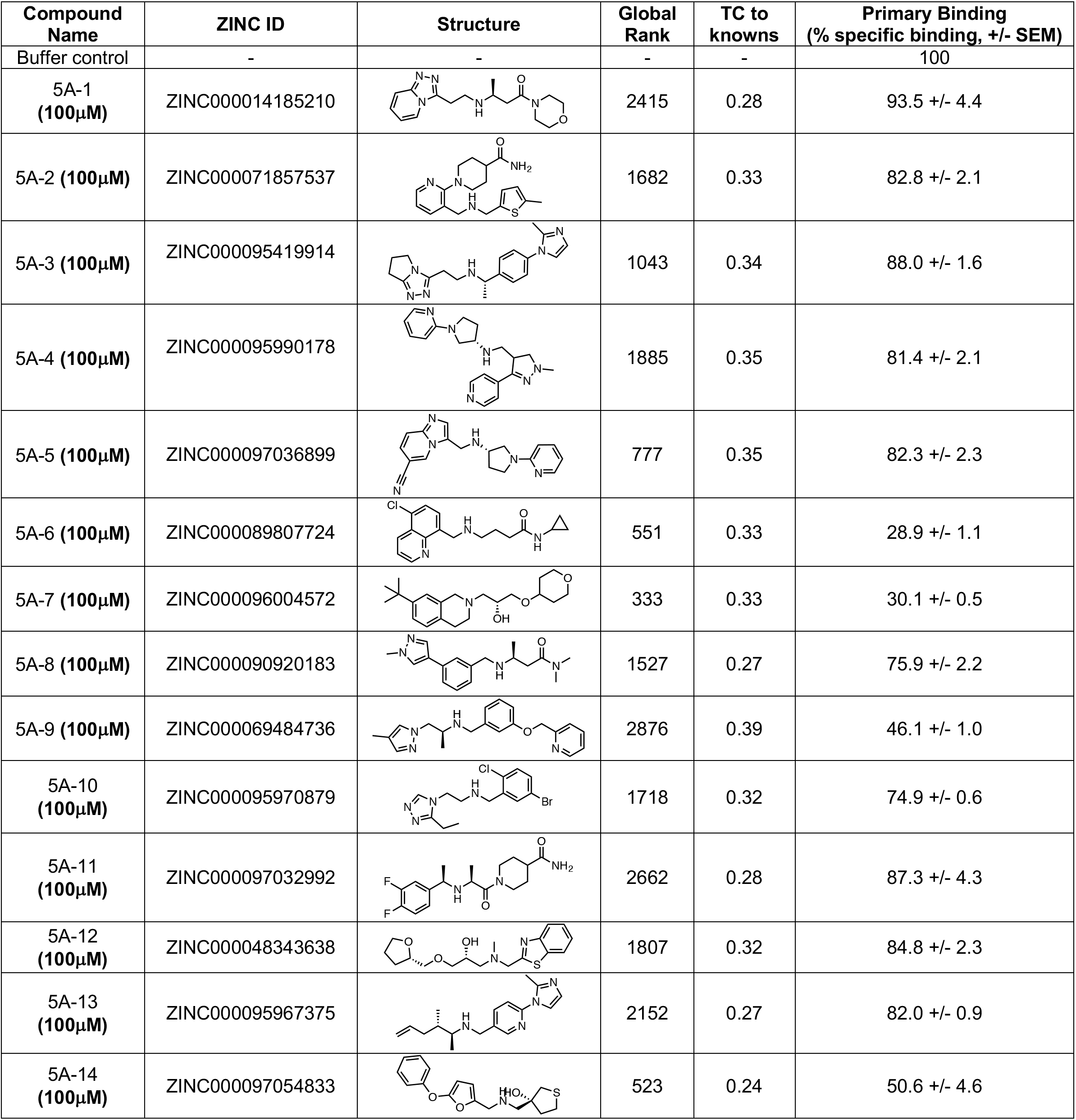
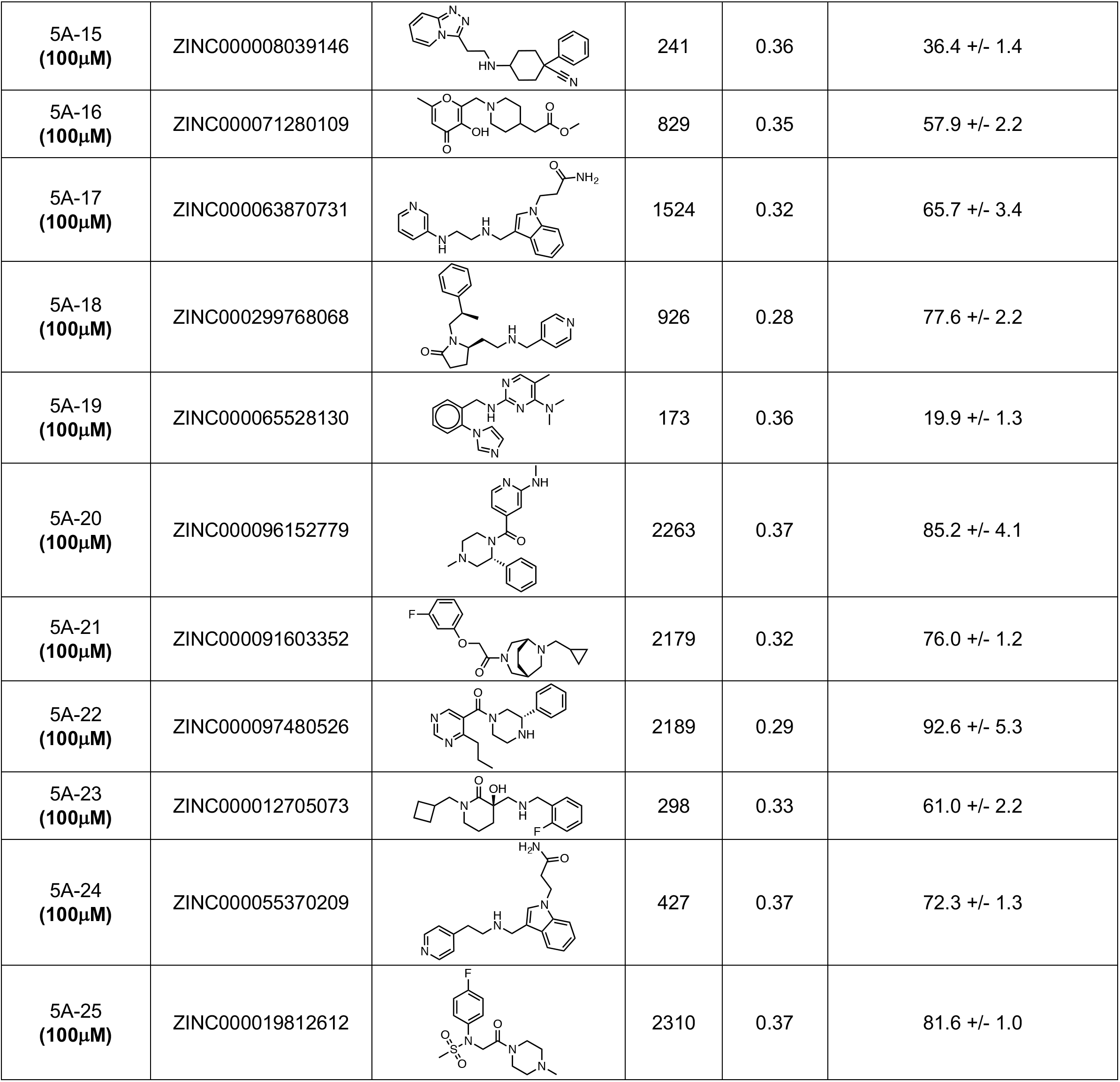
Docking hits tested at 5-HT_5A_ receptor.

| Compound Name | ZINC ID | Structure | Global Rank | TC to knowns | Primary Binding (% specific binding, +/- SEM) |
| --- | --- | --- | --- | --- | --- |
| Buffer control | - | - | - | - | 100 |
| 5A-1 (100μM) | ZINC000014185210 |  | 2415 | 0.28 | 93.5 +/- 4.4 |
| 5A-2 (100μM) | ZINC000071857537 |  | 1682 | 0.33 | 82.8 +/- 2.1 |
| 5A-3 (100μM) | ZINC000095419914 |  | 1043 | 0.34 | 88.0 +/- 1.6 |
| 5A-4 (100μM) | ZINC000095990178 |  | 1885 | 0.35 | 81.4 +/- 2.1 |
| 5A-5 (100μM) | ZINC000097036899 |  | 777 | 0.35 | 82.3 +/- 2.3 |
| 5A-6 (100μM) | ZINC000089807724 |  | 551 | 0.33 | 28.9 +/- 1.1 |
| 5A-7 (100μM) | ZINC000096004572 |  | 333 | 0.33 | 30.1 +/- 0.5 |
| 5A-8 (100μM) | ZINC000090920183 |  | 1527 | 0.27 | 75.9 +/- 2.2 |
| 5A-9 (100μM) | ZINC000069484736 |  | 2876 | 0.39 | 46.1 +/- 1.0 |
| 5A-10 (100μM) | ZINC000095970879 |  | 1718 | 0.32 | 74.9 +/- 0.6 |
| 5A-11 (100μM) | ZINC000097032992 |  | 2662 | 0.28 | 87.3 +/- 4.3 |
| 5A-12 (100μM) | ZINC000048343638 |  | 1807 | 0.32 | 84.8 +/- 2.3 |
| 5A-13 (100μM) | ZINC000095967375 |  | 2152 | 0.27 | 82.0 +/- 0.9 |
| 5A-14 (100μM) | ZINC000097054833 |  | 523 | 0.24 | 50.6 +/- 4.6 |
| 5A-15<br>(100μM) | ZINC000008039146 |  | 241 | 0.36 | 36.4 +/- 1.4 |
| 5A-16<br>(100μM) | ZINC000071280109 |  | 829 | 0.35 | 57.9 +/- 2.2 |
| 5A-17<br>(100μM) | ZINC000063870731 |  | 1524 | 0.32 | 65.7 +/- 3.4 |
| 5A-18<br>(100μM) | ZINC000299768068 |  | 926 | 0.28 | 77.6 +/- 2.2 |
| 5A-19<br>(100μM) | ZINC000065528130 |  | 173 | 0.36 | 19.9 +/- 1.3 |
| 5A-20<br>(100μM) | ZINC000096152779 |  | 2263 | 0.37 | 85.2 +/- 4.1 |
| 5A-21<br>(100μM) | ZINC000091603352 |  | 2179 | 0.32 | 76.0 +/- 1.2 |
| 5A-22<br>(100μM) | ZINC000097480526 |  | 2189 | 0.29 | 92.6 +/- 5.3 |
| 5A-23<br>(100μM) | ZINC000012705073 |  | 298 | 0.33 | 61.0 +/- 2.2 |
| 5A-24<br>(100μM) | ZINC000055370209 |  | 427 | 0.37 | 72.3 +/- 1.3 |
| 5A-25<br>(100μM) | ZINC000019812612 |  | 2310 | 0.37 | 81.6 +/- 1.0 |

**Table S2:**
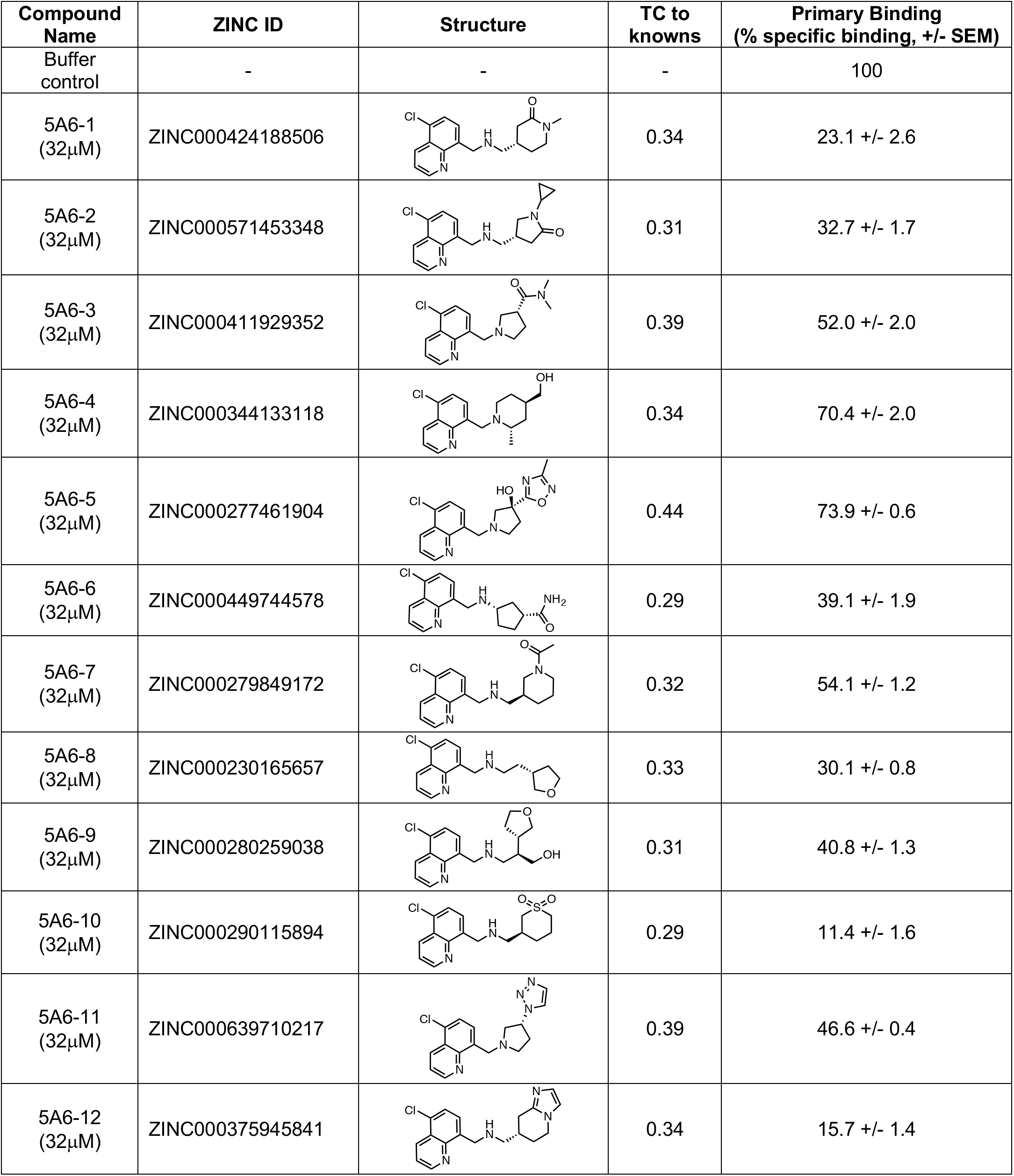
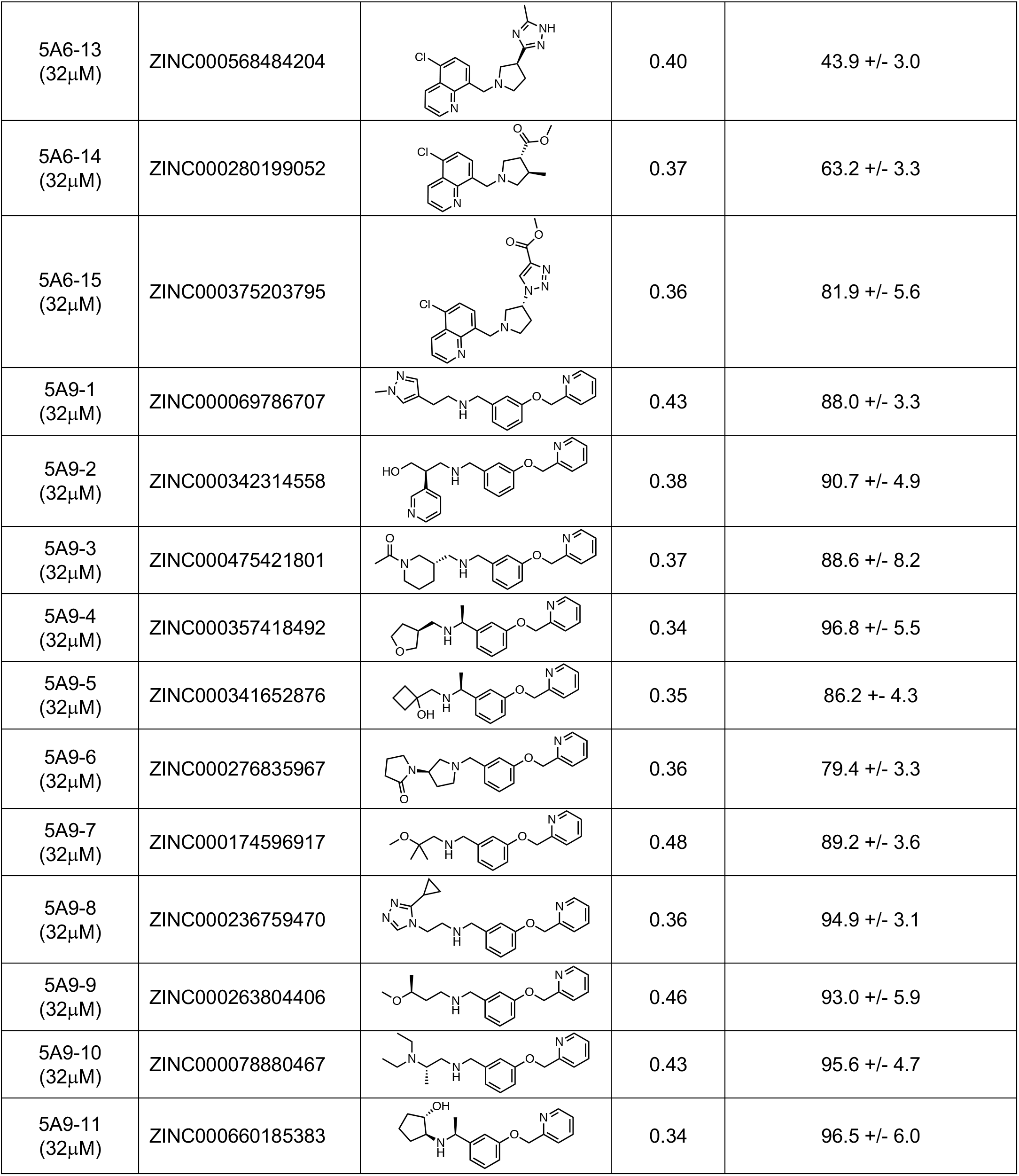
Round 1 analogs of compounds 5A-6 and 5A-9 tested at 5-HT_5A_ receptor.

| Compound Name | ZINC ID | Structure | TC to knowns | Primary Binding (% specific binding, +/- SEM) |
| --- | --- | --- | --- | --- |
| Buffer control | - | - | - | 100 |
| 5A6-1 (32μM) | ZINC000424188506 |  | 0.34 | 23.1 +/- 2.6 |
| 5A6-2 (32μM) | ZINC000571453348 |  | 0.31 | 32.7 +/- 1.7 |
| 5A6-3 (32μM) | ZINC000411929352 |  | 0.39 | 52.0 +/- 2.0 |
| 5A6-4 (32μM) | ZINC000344133118 |  | 0.34 | 70.4 +/- 2.0 |
| 5A6-5 (32μM) | ZINC000277461904 |  | 0.44 | 73.9 +/- 0.6 |
| 5A6-6 (32μM) | ZINC000449744578 |  | 0.29 | 39.1 +/- 1.9 |
| 5A6-7 (32μM) | ZINC000279849172 |  | 0.32 | 54.1 +/- 1.2 |
| 5A6-8 (32μM) | ZINC000230165657 |  | 0.33 | 30.1 +/- 0.8 |
| 5A6-9 (32μM) | ZINC000280259038 |  | 0.31 | 40.8 +/- 1.3 |
| 5A6-10 (32μM) | ZINC000290115894 |  | 0.29 | 11.4 +/- 1.6 |
| 5A6-11 (32μM) | ZINC000639710217 |  | 0.39 | 46.6 +/- 0.4 |
| 5A6-12 (32μM) | ZINC000375945841 |  | 0.34 | 15.7 +/- 1.4 |
| 5A6-13<br>(32μM) | ZINC000568484204 |  | 0.40 | 43.9 +/- 3.0 |
| 5A6-14<br>(32μM) | ZINC000280199052 |  | 0.37 | 63.2 +/- 3.3 |
| 5A6-15<br>(32μM) | ZINC000375203795 |  | 0.36 | 81.9 +/- 5.6 |
| 5A9-1<br>(32μM) | ZINC000069786707 |  | 0.43 | 88.0 +/- 3.3 |
| 5A9-2<br>(32μM) | ZINC000342314558 |  | 0.38 | 90.7 +/- 4.9 |
| 5A9-3<br>(32μM) | ZINC000475421801 |  | 0.37 | 88.6 +/- 8.2 |
| 5A9-4<br>(32μM) | ZINC000357418492 |  | 0.34 | 96.8 +/- 5.5 |
| 5A9-5<br>(32μM) | ZINC000341652876 |  | 0.35 | 86.2 +/- 4.3 |
| 5A9-6<br>(32μM) | ZINC000276835967 |  | 0.36 | 79.4 +/- 3.3 |
| 5A9-7<br>(32μM) | ZINC000174596917 |  | 0.48 | 89.2 +/- 3.6 |
| 5A9-8<br>(32μM) | ZINC000236759470 |  | 0.36 | 94.9 +/- 3.1 |
| 5A9-9<br>(32μM) | ZINC000263804406 |  | 0.46 | 93.0 +/- 5.9 |
| 5A9-10<br>(32μM) | ZINC000078880467 |  | 0.43 | 95.6 +/- 4.7 |
| 5A9-11<br>(32μM) | ZINC000660185383 |  | 0.34 | 96.5 +/- 6.0 |

**Table S3:**
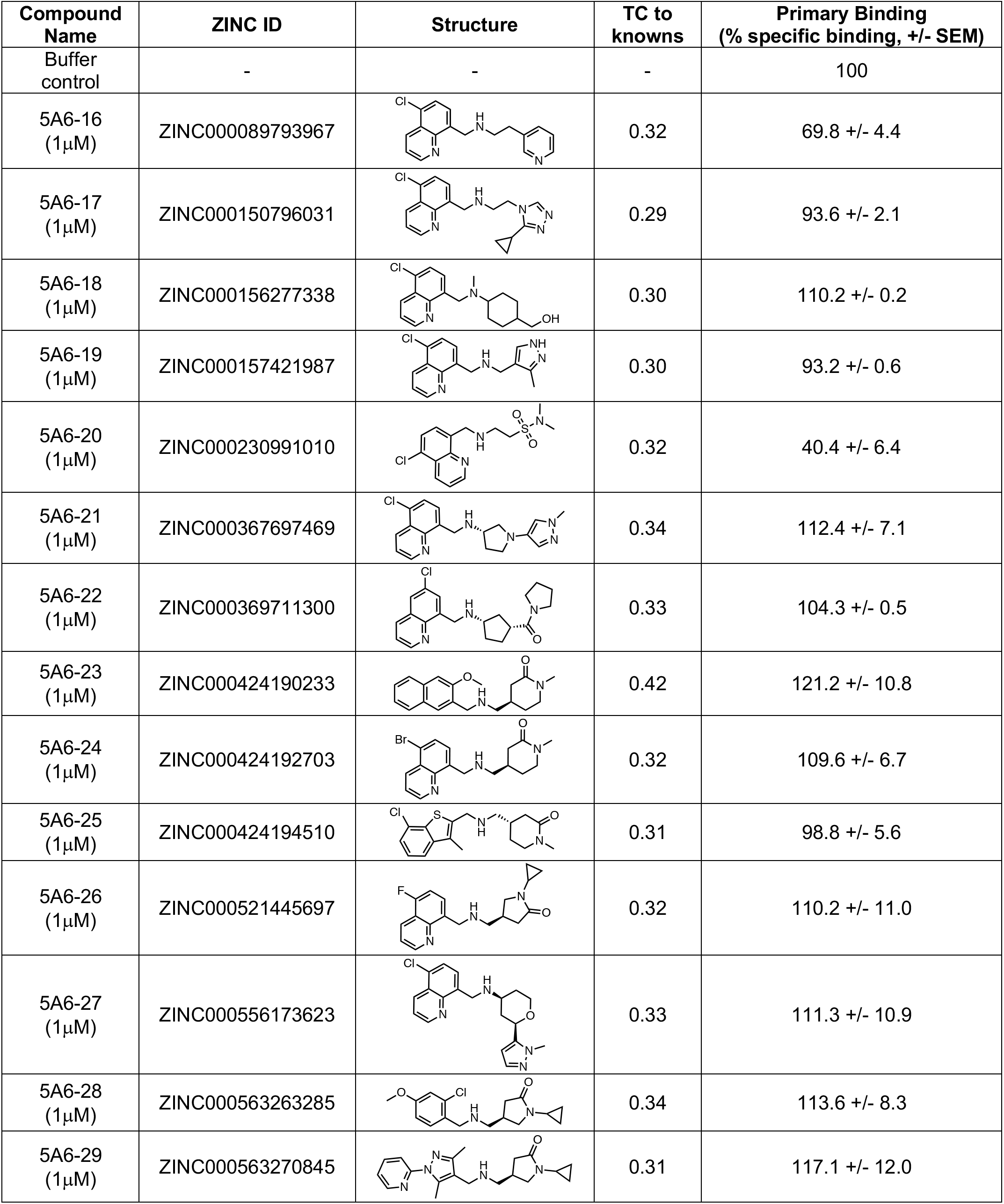
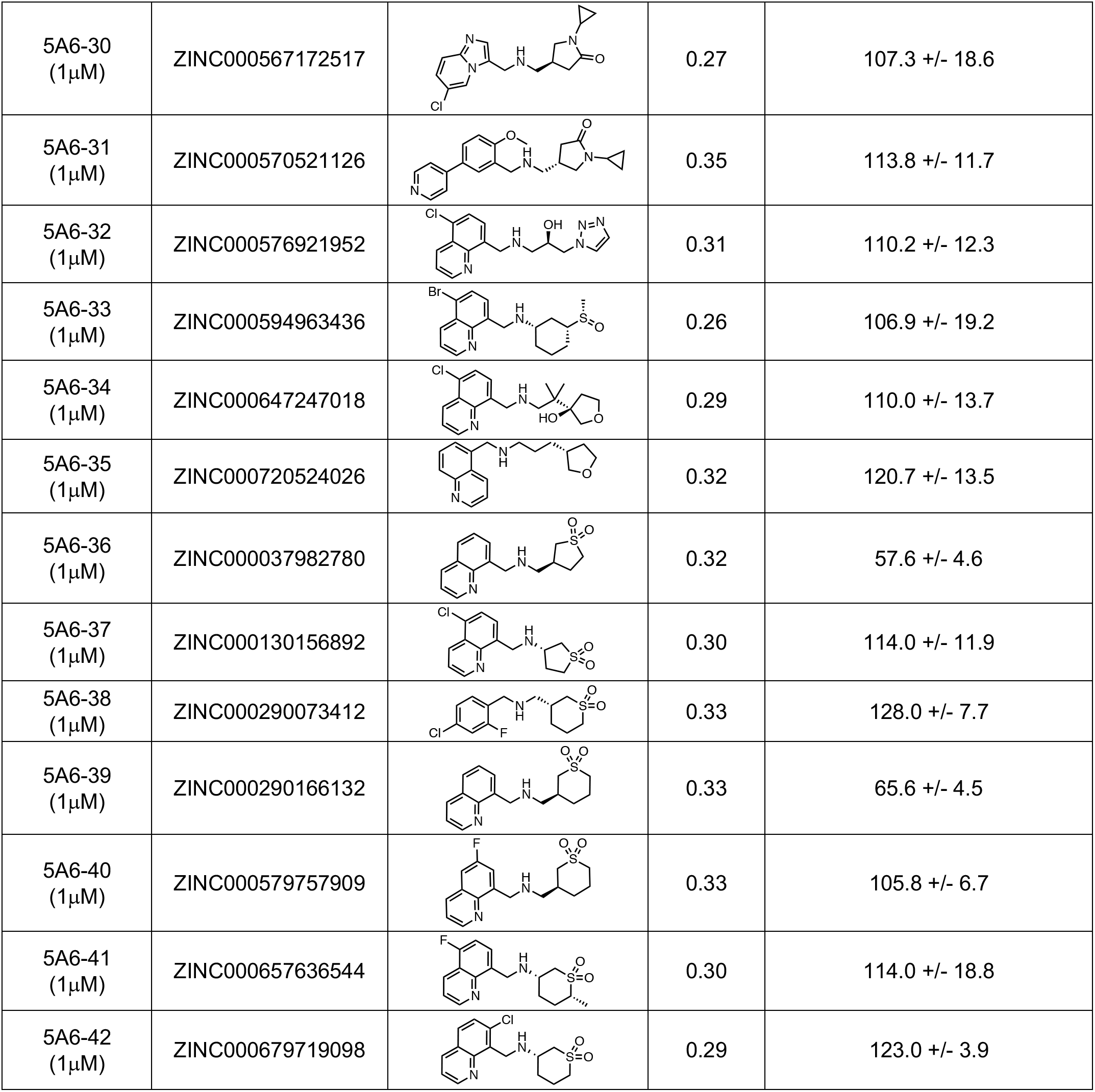
Round 2 analogs tested at 5-HT_5A_ receptor.

| Compound Name | ZINC ID | Structure | TC to knowns | Primary Binding (% specific binding, +/- SEM) |
| --- | --- | --- | --- | --- |
| Buffer control | - | - | - | 100 |
| 5A6-16 (1μM) | ZINC000089793967 |  | 0.32 | 69.8 +/- 4.4 |
| 5A6-17 (1μM) | ZINC000150796031 |  | 0.29 | 93.6 +/- 2.1 |
| 5A6-18 (1μM) | ZINC000156277338 |  | 0.30 | 110.2 +/- 0.2 |
| 5A6-19 (1μM) | ZINC000157421987 |  | 0.30 | 93.2 +/- 0.6 |
| 5A6-20 (1μM) | ZINC000230991010 |  | 0.32 | 40.4 +/- 6.4 |
| 5A6-21 (1μM) | ZINC000367697469 |  | 0.34 | 112.4 +/- 7.1 |
| 5A6-22 (1μM) | ZINC000369711300 |  | 0.33 | 104.3 +/- 0.5 |
| 5A6-23 (1μM) | ZINC000424190233 |  | 0.42 | 121.2 +/- 10.8 |
| 5A6-24 (1μM) | ZINC000424192703 |  | 0.32 | 109.6 +/- 6.7 |
| 5A6-25 (1μM) | ZINC000424194510 |  | 0.31 | 98.8 +/- 5.6 |
| 5A6-26 (1μM) | ZINC000521445697 |  | 0.32 | 110.2 +/- 11.0 |
| 5A6-27 (1μM) | ZINC000556173623 |  | 0.33 | 111.3 +/- 10.9 |
| 5A6-28 (1μM) | ZINC000563263285 |  | 0.34 | 113.6 +/- 8.3 |
| 5A6-29 (1μM) | ZINC000563270845 |  | 0.31 | 117.1 +/- 12.0 |
| 5A6-30<br>(1μM) | ZINC000567172517 | | 0.27 | 107.3 +/- 18.6 |
| 5A6-31<br>(1μM) | ZINC000570521126 | | 0.35 | 113.8 +/- 11.7 |
| 5A6-32<br>(1μM) | ZINC000576921952 | | 0.31 | 110.2 +/- 12.3 |
| 5A6-33<br>(1μM) | ZINC000594963436 | | 0.26 | 106.9 +/- 19.2 |
| 5A6-34<br>(1μM) | ZINC000647247018 | | 0.29 | 110.0 +/- 13.7 |
| 5A6-35<br>(1μM) | ZINC000720524026 | | 0.32 | 120.7 +/- 13.5 |
| 5A6-36<br>(1μM) | ZINC000037982780 | | 0.32 | 57.6 +/- 4.6 |
| 5A6-37<br>(1μM) | ZINC000130156892 | | 0.30 | 114.0 +/- 11.9 |
| 5A6-38<br>(1μM) | ZINC000290073412 | | 0.33 | 128.0 +/- 7.7 |
| 5A6-39<br>(1μM) | ZINC000290166132 | | 0.33 | 65.6 +/- 4.5 |
| 5A6-40<br>(1μM) | ZINC000579757909 | | 0.33 | 105.8 +/- 6.7 |
| 5A6-41<br>(1μM) | ZINC000657636544 | | 0.30 | 114.0 +/- 18.8 |
| 5A6-42<br>(1μM) | ZINC000679719098 | | 0.29 | 123.0 +/- 3.9 |

**Table S4:**
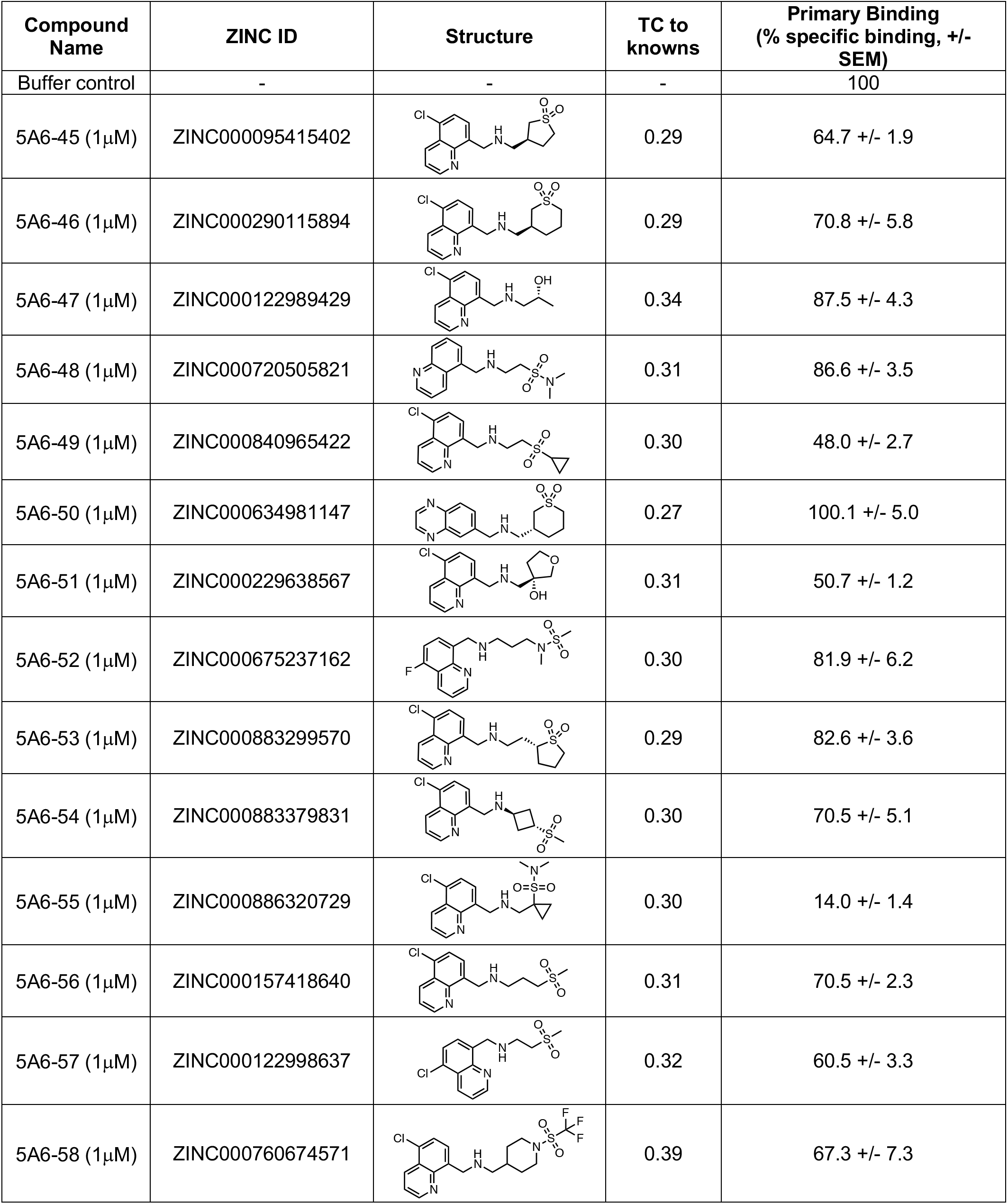
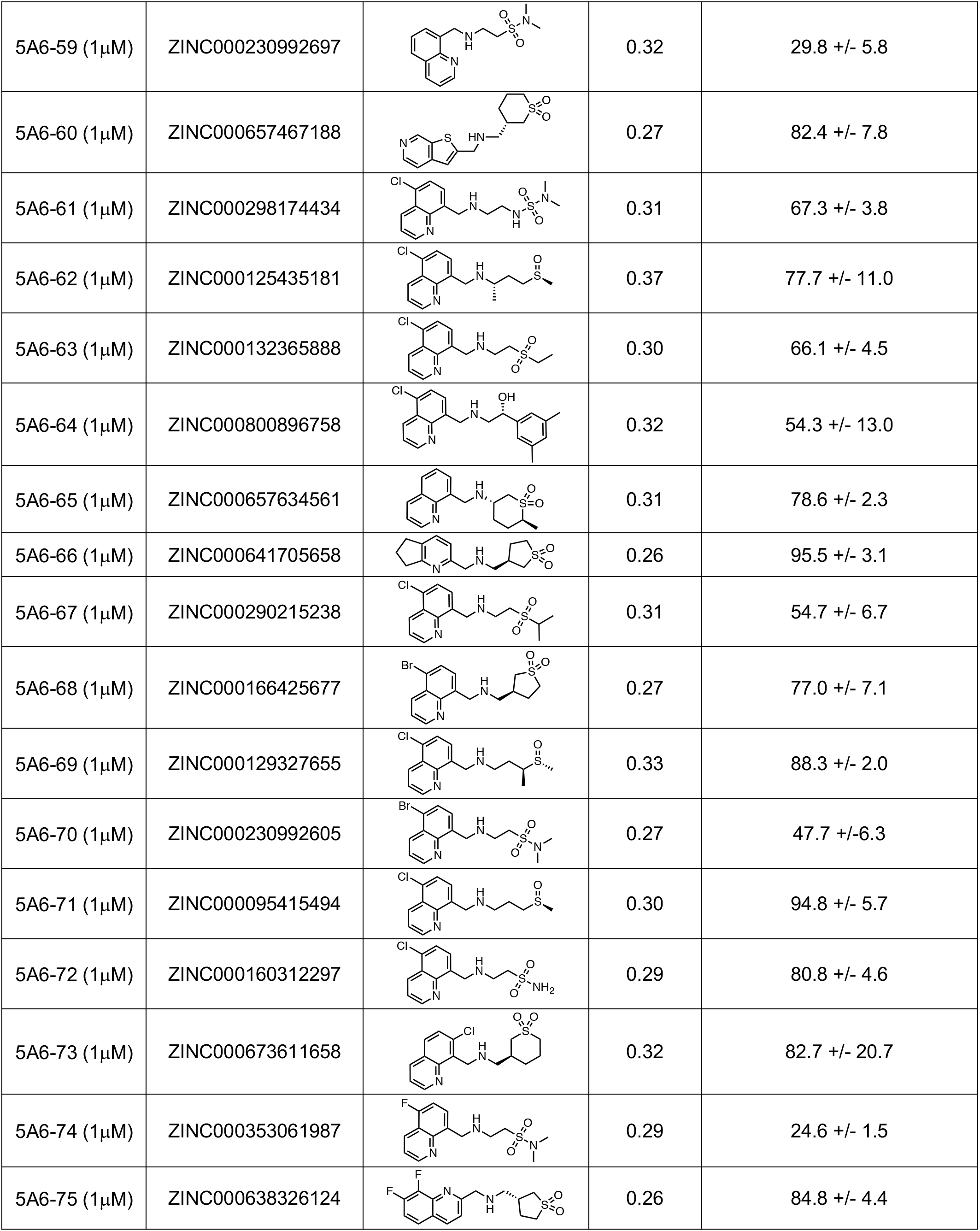
Round 3 analogs tested at 5-HT_5A_ receptor.

| Compound Name | ZINC ID | Structure | TC to knowns | Primary Binding (% specific binding, +/- SEM) |
| --- | --- | --- | --- | --- |
| Buffer control | - | - | - | 100 |
| 5A6-45 (1μM) | ZINC000095415402 |  | 0.29 | 64.7 +/- 1.9 |
| 5A6-46 (1μM) | ZINC000290115894 |  | 0.29 | 70.8 +/- 5.8 |
| 5A6-47 (1μM) | ZINC000122989429 |  | 0.34 | 87.5 +/- 4.3 |
| 5A6-48 (1μM) | ZINC000720505821 |  | 0.31 | 86.6 +/- 3.5 |
| 5A6-49 (1μM) | ZINC000840965422 |  | 0.30 | 48.0 +/- 2.7 |
| 5A6-50 (1μM) | ZINC000634981147 |  | 0.27 | 100.1 +/- 5.0 |
| 5A6-51 (1μM) | ZINC000229638567 |  | 0.31 | 50.7 +/- 1.2 |
| 5A6-52 (1μM) | ZINC000675237162 |  | 0.30 | 81.9 +/- 6.2 |
| 5A6-53 (1μM) | ZINC000883299570 |  | 0.29 | 82.6 +/- 3.6 |
| 5A6-54 (1μM) | ZINC000883379831 |  | 0.30 | 70.5 +/- 5.1 |
| 5A6-55 (1μM) | ZINC000886320729 |  | 0.30 | 14.0 +/- 1.4 |
| 5A6-56 (1μM) | ZINC000157418640 |  | 0.31 | 70.5 +/- 2.3 |
| 5A6-57 (1μM) | ZINC000122998637 |  | 0.32 | 60.5 +/- 3.3 |
| 5A6-58 (1μM) | ZINC000760674571 |  | 0.39 | 67.3 +/- 7.3 |
| 5A6-59 (1μM) | ZINC000230992697 |  | 0.32 | 29.8 +/- 5.8 |
| 5A6-60 (1μM) | ZINC000657467188 |  | 0.27 | 82.4 +/- 7.8 |
| 5A6-61 (1μM) | ZINC000298174434 |  | 0.31 | 67.3 +/- 3.8 |
| 5A6-62 (1μM) | ZINC000125435181 |  | 0.37 | 77.7 +/- 11.0 |
| 5A6-63 (1μM) | ZINC000132365888 |  | 0.30 | 66.1 +/- 4.5 |
| 5A6-64 (1μM) | ZINC000800896758 |  | 0.32 | 54.3 +/- 13.0 |
| 5A6-65 (1μM) | ZINC000657634561 |  | 0.31 | 78.6 +/- 2.3 |
| 5A6-66 (1μM) | ZINC000641705658 |  | 0.26 | 95.5 +/- 3.1 |
| 5A6-67 (1μM) | ZINC000290215238 |  | 0.31 | 54.7 +/- 6.7 |
| 5A6-68 (1μM) | ZINC000166425677 |  | 0.27 | 77.0 +/- 7.1 |
| 5A6-69 (1μM) | ZINC000129327655 |  | 0.33 | 88.3 +/- 2.0 |
| 5A6-70 (1μM) | ZINC000230992605 |  | 0.27 | 47.7 +/- 6.3 |
| 5A6-71 (1μM) | ZINC000095415494 |  | 0.30 | 94.8 +/- 5.7 |
| 5A6-72 (1μM) | ZINC000160312297 |  | 0.29 | 80.8 +/- 4.6 |
| 5A6-73 (1μM) | ZINC000673611658 |  | 0.32 | 82.7 +/- 20.7 |
| 5A6-74 (1μM) | ZINC000353061987 |  | 0.29 | 24.6 +/- 1.5 |
| 5A6-75 (1μM) | ZINC000638326124 |  | 0.26 | 84.8 +/- 4.4 |

**Table S5:**
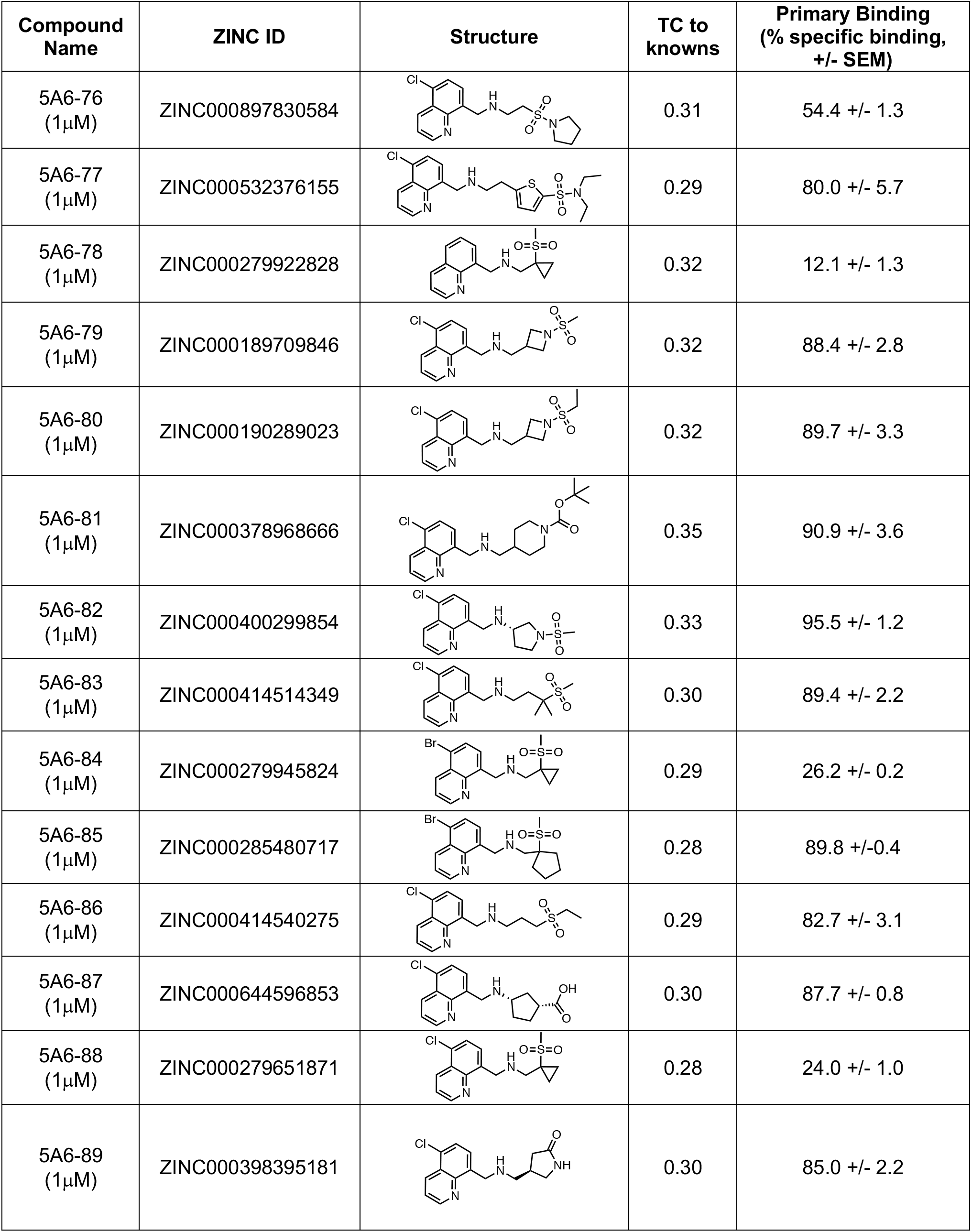
Round 4 analogs tested at 5-HT_5A_ receptor.

| Compound Name | ZINC ID | Structure | TC to knowns | Primary Binding (% specific binding, +/- SEM) |
| --- | --- | --- | --- | --- |
| 5A6-76 (1μM) | ZINC000897830584 |  | 0.31 | 54.4 +/- 1.3 |
| 5A6-77 (1μM) | ZINC000532376155 |  | 0.29 | 80.0 +/- 5.7 |
| 5A6-78 (1μM) | ZINC000279922828 |  | 0.32 | 12.1 +/- 1.3 |
| 5A6-79 (1μM) | ZINC000189709846 |  | 0.32 | 88.4 +/- 2.8 |
| 5A6-80 (1μM) | ZINC000190289023 |  | 0.32 | 89.7 +/- 3.3 |
| 5A6-81 (1μM) | ZINC000378968666 |  | 0.35 | 90.9 +/- 3.6 |
| 5A6-82 (1μM) | ZINC000400299854 |  | 0.33 | 95.5 +/- 1.2 |
| 5A6-83 (1μM) | ZINC000414514349 |  | 0.30 | 89.4 +/- 2.2 |
| 5A6-84 (1μM) | ZINC000279945824 |  | 0.29 | 26.2 +/- 0.2 |
| 5A6-85 (1μM) | ZINC000285480717 |  | 0.28 | 89.8 +/- 0.4 |
| 5A6-86 (1μM) | ZINC000414540275 |  | 0.29 | 82.7 +/- 3.1 |
| 5A6-87 (1μM) | ZINC000644596853 |  | 0.30 | 87.7 +/- 0.8 |
| 5A6-88 (1μM) | ZINC000279651871 |  | 0.28 | 24.0 +/- 1.0 |
| 5A6-89 (1μM) | ZINC000398395181 |  | 0.30 | 85.0 +/- 2.2 |

